# Fetal sex influences molecular regulation of human placental vascular development across gestation

**DOI:** 10.64898/2026.09.13.750572

**Authors:** Madalena Corado, Luana Ferreira Afonso, Sofia P. Agostinho, Irina S. Moreira, Nícia Rosário-Ferreira, Vanessa Coelho-Santos

## Abstract

Males experience elevated mortality and morbidity during the perinatal period and infancy; however, the biological origins of this vulnerability remain unclear. Emerging evidence suggests that sex-dependent placental development may contribute to differences in fetal growth and outcomes. Vascular networks are central to placental function, established early in the first trimester to support growth and mitigate harmful exposures, and maturing by term to sustain late fetal development. Although these processes are tightly regulated by gene-expression programmes, the role of sex-specific vascular signalling in placental development remains poorly understood.

To address this, we analysed raw, publicly available RNA-sequencing transcriptomics data from first trimester and term placental samples. We conducted parallel differential gene expression and functional enrichment analyses across the datasets to ensure direct comparability and to disentangle the specific contribution of sex chromosomes.

We identified multiple differentially expressed genes and enriched pathways between male and female placentas. In the first trimester, male placentas exhibited significant enrichment of vascular signalling pathways, particularly Notch signalling, cell adhesion, and migration, consistent with heightened proliferative and remodelling activity. At the end of gestation, sex differences remained detectable; however differentially expressed genes was clearly sex-chromosome driven, with autosomal single-gene sex differences and autosomal-sex-specific pathways markedly reduced. This reflected a mature placental vascular system with minimal sex-based differences outside of the sex chromosome genes.

These results reveal gestational stage-dependent sex differences in placental vascular development, with early male fetuses exhibiting increased angiogenic activity and immature vascular signalling, potentially contributing to accelerated growth and increased vulnerability in male neonates.

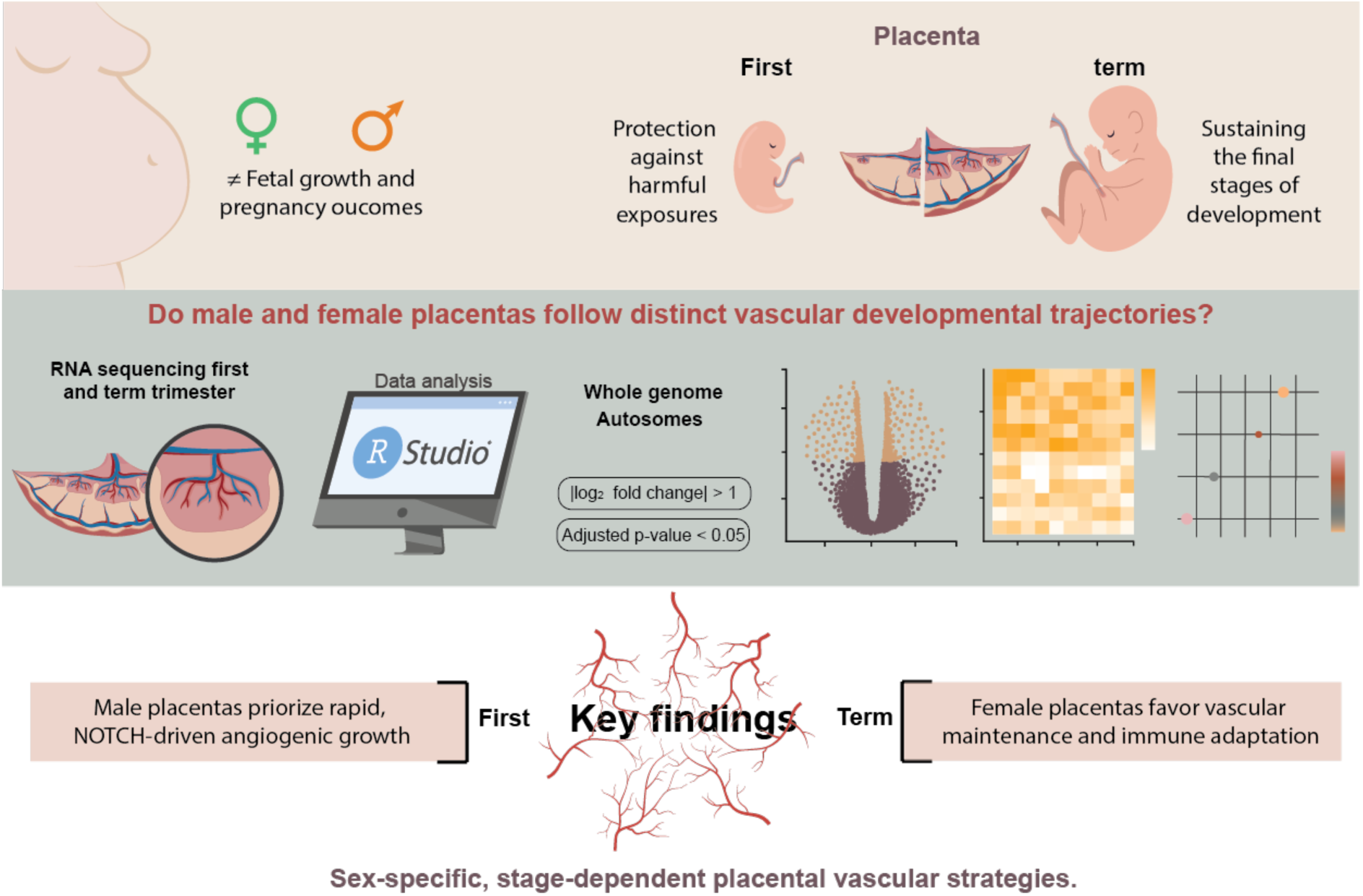

## Introduction

The placenta is a remarkable organ that forms the interface between mother and fetus, essential for supporting fetal growth and health. Its circulation is comprises of two separate blood supplies, maternal and fetal, that enable the delivery of oxygen and nutrients, the removal of waste products, and the regulation of immune interactions throughout gestation ^1^. Beyond nutrient and gas exchange, the placenta also protects the fetus from harmful exposures to environmental toxins and physical insults. However, this protection is not absolute, and studies have demonstrated that fetuses can still be exposed to drugs and other substances due to the placenta’s selective permeability ^2–4^.

The placental barrier resides on the surface of placental villi, highly vascularized finger-like projections extending into the maternal blood space. Each villus contains a core of mesenchymal tissue with fetal capillaries, surrounded by two layers of epithelial trophoblasts, the primary placental cells ^5^. The placental barrier itself comprises three distinct layers: the syncytiotrophoblast, the cytotrophoblast, and a fetal capillary endothelium. The first two layers are composed of trophoblasts, and constitute the outer and inner layers, respectively, that are in contact with the maternal blood supply. Meanwhile, the fetal capillary endothelium lines the fetal blood vessels within the villi ^6^. Malformations of the villous tree can result in an underdeveloped vascular network and associated pregnancy complications, including miscarriage, preeclampsia, stillbirth, or preterm labour ^7–10^. Additionally, it can result to fetal growth restriction (FGR) ^11^, contributing to stunted fetal development and growth, and an elevated risk of postnatal neurological disorders, such as autism spectrum disorder ^12–14^.

Central to fetal development is the establishment and expansion of a robust placental vascular system with profound implications for health and development both in utero and beyond. The placenta begins to develop from the first weeks of gestation, from a simple mesenchymal villi that progressively evolves into a placental villi, more complex and irrigated ^15^. Throughout gestation, the placental vascular network undergoes continuous remodelling, driven by two essential processes: vasculogenesis, the formation of new blood vessels, and angiogenesis, the growth of vessels from existing ones. During early gestation, vasculogenesis establishes the primary vascular network ^16^, which is subsequently expanded during the first and second trimesters via angiogenesis. By the end of the second trimester, this vascular expansion results in an accentuated increase in capillary length, volume, and surface area, driven by endothelial cell proliferation, culminating in an extensive and efficient capillary network at term ^17^. Supporting the expansion and remodelling of the placental vascular network are epithelial trophoblasts, which directly connect with the maternal vascular network and tightly regulate several signalling pathways. Differentiated trophoblasts express vascular endothelial growth factors (*VEGF*) 1 and 2, key drivers of blood vessel formation ^18^. Defective VEGF signalling has been associated with impaired blood flow and insufficient nutrient delivery to the fetus ^19^. Furthermore, trophoblasts express placental growth factor (*PlGF*), a pro-angiogenic member of the *VEGF* family that enhances angiogenesis by activating *VEGF* signalling pathways and amplifying *VEGF* activity ^18^.

Fetal sex is a significant determinant of placental development and function, growth trajectories, and pregnancy outcomes from early gestation. As early as 8–12 weeks gestation, female fetuses are, on average, smaller than males, a difference that persists throughout pregnancy ^20^. These sex differences extend well beyond fetal size, encompassing divergent placental and fetal responses to biological signalling molecules ^21^, maternal and environmental factors ^21^, and even drug exposures ^22^. Despite increased growth, males experience consistently higher rates of morbidity and mortality during the perinatal period and infancy ^23,24^, and are at increased risk of spontaneous abortion during the first trimester ^25^. This male vulnerability is hypothesized to reflect evolutionary differences operating across gestation and the perinatal period; however, the precise mechanisms underlying the female survival advantage remain unclear. Accumulating evidence suggests that these sex disparities are inherently multifactorial, arising from the interplay of differential sex steroid exposure, X-chromosome gene dosage and regulation, and sex-specific placental adaptive responses ^26–28^.

The placental vasculature is central to sustaining fetal growth through efficient nutrient, gas, and waste exchange, yet sex-specific differences in vascular signalling across gestation remain poorly defined. Notably, pregnancies carrying female fetuses, in both healthy and preeclamptic pregnancies, tend to exhibit higher levels of the angiogenic markers soluble Fms-like kinase 1 (*sFlt-1*) and elevated sFlt-1/ *PlGF* ratios compared to those carrying male fetuses, suggesting distinct angiogenic regulatory environments ^29^. Furthermore, during the first gestational trimester, *PIGF* levels are lower in male fetuses than in female fetuses, but by the second trimester, *PIGF* levels rise sharply in male while remaining relatively stable in females ^30^. Together, these observations suggest fundamentally divergent angiogenic trajectories according to fetal sex, with potential implications for placental resilience and fetal survival. In this study, we investigate sex-specific differences in vascular-related placental gene expression during early and late gestation to define molecular pathways potentially contributing to differential fetal vulnerability.

## Methods

All data analyses and visualizations, described in the following subsection were performed in R (version 4.5.1). All codes used for RNA-seq preprocessing, differential expression analysis, and data visualization for first trimester and term placenta samples, with full datasets and an autosomal-only version, are publicly available at https://github.com/MoreiraLAB/FeSPlaVT.git and the detailed description of package versions are presented in Supplementary Table 1.

### Dataset and sample characteristics

To investigate sex-based differences in vascular genes and pathways during the first trimester and term placental development, we analysed bulk RNA-Sequencing raw count data from two independent transcriptomic datasets. The first dataset was obtained from a SMAART cohort (GEO accession: GSE215421) ^31^. The second dataset was kindly provided by Dr. Melissa A. Wilson, Senior Investigator at the National Human Genome Research Institute, National Institutes of Health, and was previously described in detail ^26^.

First trimester samples (n=141) consisted of chorionic villus tissue collected between 10-14 weeks of gestation from singleton pregnancies. Pregnancies were conceived spontaneously (74) or through fertility treatments, including *in vitro* fertilization (IVF) (34) and non-*in vitro* fertilization (NIFT) (33). The cohort included 72 male and 69 female fetuses. All samples were collected from individuals of Caucasian or biracial Caucasian/Asian descent.

The term dataset comprised chorionic placental tissue collected after cesarean delivery at ≧ 36 weeks of gestation (n=60). The cohort included 30 male and 30 female fetuses. Fifty-eight pregnancies were spontaneous; one resulted from IVF and one from NIFT. The cohort encompassed several self-reported ethnicities, including Caucasian, Asian, Black, and Hispanic. Samples were sequenced in two batches: 12 males and 12 females in the first batch, and 18 males and 18 females in the second.

### RNA-seq preprocessing and filtering

Parallel preprocessing workflows were applied to the raw count matrices of both datasets. For the first trimester dataset, two samples were excluded due to inconsistencies in sample annotation across files, resulting in a final dataset of 139 samples. Prior to downstream analysis, 50 genes corresponding to ribosomal RNA and hemoglobin transcripts were removed. To enhance model stability and reduce noise, genes with low counts were filtered, retaining only those with a total count of ≥ 10 reads summed across all samples. This filtering step reduced the gene set from 39,326 to 34,197 genes.

For the term dataset, one biological sample (YPOPS0007M) was excluded due to the absence of a technical replicate in the count matrix, resulting in 59 biological term placenta samples with a total of 118 technical replicates. Following initial filtering of 57,133 genes, 29 corresponding to ribosomal RNA and hemoglobin transcripts were removed. Technical replicates were subsequently collapsed using the collapseReplicates function from the DESeq2 package, preventing duplicate sample entries in downstream analyses. Low-count gene filtering was similarly applied, retaining genes with total read counts of ≧ 10 across samples, which reduced the feature set from 57,104 to 38,144 genes. This measure reduces sparsity and improves model stability, consistent with established recommendations for DESeq2.

For both first trimester and term placenta RNA-seq datasets, we additionally generated autosome-only count matrices by excluding all genes located on sex chromosomes (X and Y). In the first trimester dataset, RefSeq accession numbers (ChrAcc) from the NCBI annotation file (Human.GRCh38.p13.annot.tsv.gz) were used to identify the sex chromosome genes, with NC_000023.11 indicating X chromosome genes and NC_000024.10 indicating Y chromosome genes. In the term dataset, genes assigned to sex chromosomes were identified through the “chr” information (“chrx”, “chry”) provided in the raw counts file (placenta_batch1and2_geneCounts.tsv). Subsequent analyses were performed in parallel on both the whole-genome datasets and the corresponding autosome-only matrices to assess whether autosomal regulation differs from direct sex-chromosome dosage.

### Dataset characterization and DESeq2 model design

To design robust differential expression models, we systematically evaluated the distribution of biological and technical variables across both datasets. This initial characterization was essential for identifying key confounders, technical artifacts, and biological covariates critical for accurate DESeq2 model specification.

To specifically guide model selection, we employed Variance Partition analysis to quantitatively evaluate the contribution of technical (e.g. batch) and biological (e.g. ancestry) covariates to overall gene expression variance. This approach fits linear mixed models to the variance-stabilized expression matrix, estimating the proportion of variance attributable to each covariate per gene.

Complementing this quantitative assessment, Principal Component Analysis (PCA) was conducted to visualize the major axes of transcriptional variability. PCA was performed independently for the first trimester and term datasets utilizing variance-stabilized expression values (generated with the *vst()* function from DESeq2, with blind=TRUE’ to ensure unbiased estimation). Principal components were derived using the DESeq2’s *plotPCA()* function, with sample covariates supplied via the *intgroup* argument to highlight factors of interest (e.g. sex, batch). This analysis was performed on both the full gene set and the top 500 most variable genes to capture global data structure. No additional outliers were removed based on PCA or sample-to-sample distances.

### Differential expression analysis and visualization

Differential gene expression analysis was performed using the DESeq2 package ^32^, with female samples set as the reference category for fetal sex comparisons. DESeqDataSet objects were constructed using the DESeq2 function *DESeqDataSetFromMatrix()*, providing the filtered raw count matrix, corresponding sample metadata and the experimental design formula. Hypothesis testing for differential expression was performed using the Wald test, as implemented in DESeq2’s *DESeq()* function ^33^, which also estimates size factors and dispersion parameters for each gene within a negative binomial generalized linear model framework. Following model fitting, differential expression results were extracted for contrasts of interest using the *results()* function. To improve effect size estimation, log2 fold changes were shrunk using the *lfcShrink()* function with the “apeglm” method ^34^, reducing noise, particularly for lowly expressed or highly variable genes. Multiple testing correction was performed using the Benjamini-Hochberg procedure to control the false discovery rate (FDR) ^35^. Genes were considered differentially expressed if they had an adjusted p-value (FDR) < 0.05 and an absolute log2 fold change > 1. The model was fitted to test fetal sex effects for both first trimester and term placenta datasets. In the first trimester dataset, the design formula incorporated RNA-sequencing batch as a technical covariate. In the term placenta dataset, the design included batch and exome sequencing principal components. Ancestry principal components were included in the design formula of the term placenta RNA-seq differential expression analysis to adjust for population stratification and genetic heterogeneity among the samples. Volcano plots were generated using the EnhancedVolcano package, utilizing input from DESeq2 result objects, and displaying log2 fold change on the x-axis and -log10 adjusted p-value on the y-axis. Genes meeting significance and fold change cutoffs were labeled and highlighted. Heatmaps were produced using’ pheatmap based on variance-stabilized transformed expression data (obtained using DESeq2’s vst() function). The top 30 Differentially expressed genes (DEGs) ranked by adjusted p-value or all genes passing significance criteria were used for heatmap generation. Hierarchical clustering was performed on genes using correlation distance and on samples using Euclidean distance to reveal co-expression and sample grouping patterns.

### Functional Enrichment

Functional enrichment analysis was conducted using a rank-based Gene Set Enrichment Analysis (GSEA) approach. Differential expression results from DESeq2 were used to rank gene sets; specifically, the unshrunken Wald test statistic from the primary DESeq2 output was extracted and utilized as the ranking metric for each dataset’s primary comparison (male versus female fetal sex). GSEA was performed separately on two main gene set groupings: core pathway and hallmark gene sets in aggregate and chromosome location gene sets. For each group, gene sets were sourced from the Molecular Signatures Database (MSigDB package v25.1.1) ^36,37^: pathway and hallmark analyses included the Hallmark collection (H) and canonical pathway gene sets from KEGG (C2:CP:KEGG_MEDICUS), Reactome (C2:CP:REACTOME), WikiPathways (C2:CP:WIKIPATHWAYS), BioCarta (C2:CP:BIOCARTA), and the Pathway Interaction Database (C2:CP:PID) and chromosome analyses were based on gene sets derived from the C1 collection of positional gene sets. For each collection, gene sets containing 15-500 genes were retained for enrichment to balance specificity and power. Enrichment significance was assessed using the fgseaMultilevel algorithm with exact p-value estimation (eps=0.0), employing a fixed random seed to ensure reproducibility. Statistical significance was determined using the Benjamini-Hochberg FDR threshold of 0.05 across all gene sets within each collection, and normalized enrichment scores (NES) were used to compare magnitude and directionality of enrichment across gene sets.

Within each of these primary analyses, an explicit subset of gene sets was selected a priori to focus on vascular- and barrier-related terms, motivated by the central biological theme of the study. These ‘vascular and barrier’ panels, available in full in the FeSPlaVT repository, were defined using a combination of domain knowledge and systematic keywords, capturing terms directly relevant to angiogenesis, vascular development, endothelial junctions, the renin-angiotensin axis, extracellular matrix remodelling, integrin signalling, platelet function, coagulation, and tight junctions. Thus, vascular-focused enrichment results are presented as a focused subset within each primary group’s analysis, complementing the hypothesis-unbiased results with targeted interrogation. Separate enrichment analyses and visualizations are reported for global and vascular-focused panels within each gene set collection.

## Results

### Pre-processing and sample distribution quality control

The overall analysis workflow, encompassing preprocessing, quality control, differential expression modelling, and gene set enrichment analyses, is summarized in **Figure 1**. Briefly, we generated curated whole-genome and autosome-only RNA-seq count matrices for first trimester and term placentas, characterized sample structure and sources of variation (batch, sex, ancestry, clinical covariates), and subsequently applied DESeq2 and GSEA to quantify sex-biased genes and pathways. Following quality control and exclusion of misannotated or incomplete samples, the first trimester cohort comprised 139 placentas (72 male, 67 female), and the term cohort consisted of 59 placentas (30 male, 29 female). Groups were broadly balanced by fetal sex within each dataset, with sequencing performed across multiple batches (five for first trimester, two for term), as detailed in **Figures S1-S2**.

**Figure 1.**
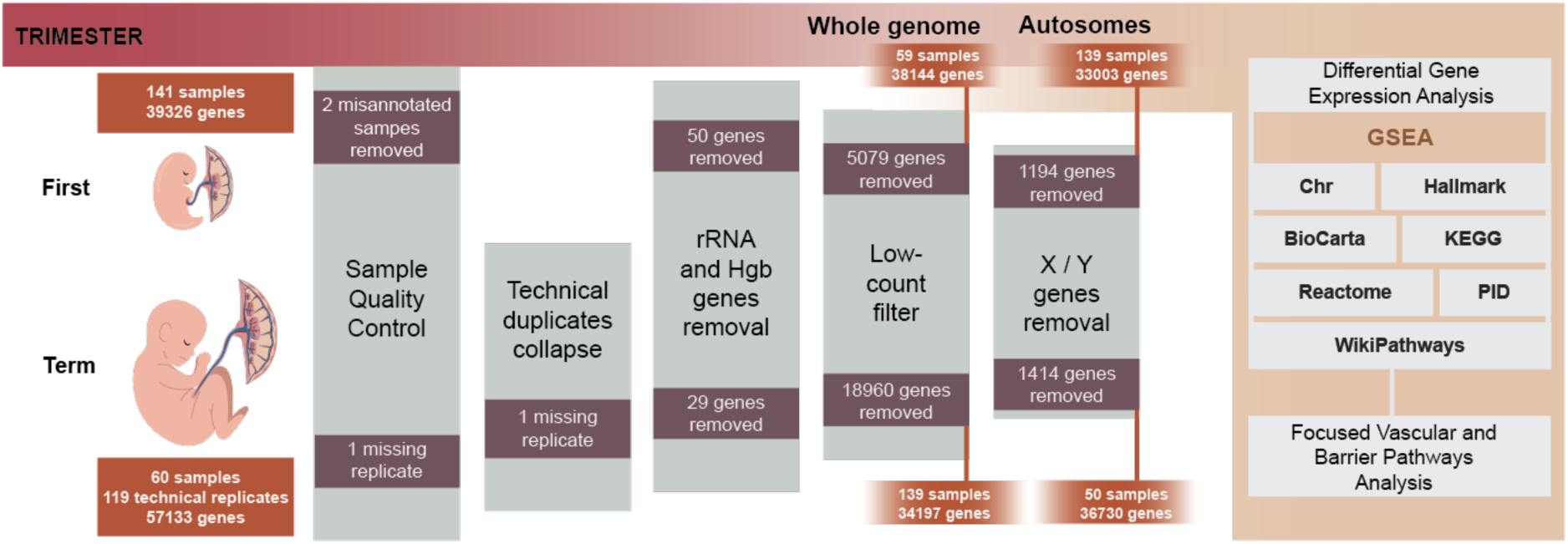
RNA-seq analysis workflow for sex-biased placental transcription. Bulk RNA-seq count matrices from first trimester and term placentas were subjected to staged quality control and filtering. Following removal of misannotated samples, ribosomal and hemoglobin genes, and low-count filtering, parallel whole-genome and autosome-only datasets were generated for each gestational stage. DESeq2 differential expression models incorporated fetal sex as the primary variable, with batch and ancestry retained as covariates based on their contribution to transcriptomic variance. Ranked differential expression statistics were then input into GSEA across multiple curated gene set collections, including pathways, hallmarks, and chromosomal regions, as well as vascular and barrier-focused modules, to identify significant findings. Original image created by a Graphic Designer (www.behance.net/TiagoFigueiredoGD).

### First trimester placenta: sample distribution

**Figure S1A** demonstrates a balanced sex ratio across spontaneous, NIFT, and IVF conceptions, with approximately equal male-to-female ratios within each mode. This finding suggests that fertility treatment type does not systematically influence sex skew. Furthermore, **Figure S1B** indicates that both male and female samples are represented across the five RNA-seq batches utilized for first trimester sequencing.

However, **Figures S1C** and **S1D** reveal a marked collinearity between sequencing batch and mode of conception. Specifically, IVF and NIFT samples are consistently found within the ART Set1 and ART Set2 groups, whereas spontaneous conceptions predominate in ART Supplemental, SexDiff Set1, and SexDiff Set2. This technical confounding, wherein batch and mode of conception are inextricably linked, necessitates consideration in model specification.

### Term placenta: sample distribution

In the term dataset, **Figure S2A** indicates an even distribution of fetal sex across the two sequencing batches. **Figure S2B** reveals that only 2 of 59 biological samples were conceived through assisted reproduction (one IVF, one IUI), with 96.6% resulting from spontaneous conception. This extreme imbalance renders mode of conception statistically non-estimable in differential expression models, as further discussed below.

**Figures S2C** and **S2D** demonstrate that mode of conception and RNA-seq batch exhibit confounding patterns comparable to those observed in the first trimester, yet with a more pronounced imbalance. **Figure S2E** reveals that birth weight displays expected sex-biased variation, with males exhibiting a higher median birth weight than females, consistent with existing literature ^38^. However, birth weight is an outcome of pregnancy reflecting downstream consequences of sex differences in gene expression, metabolism, and growth, rather than a causal determinant of placental transcriptomic differences. Consequently, incorporating such outcome variables into the model risks biasing effect estimates for the primary variable, fetal sex.

Reported race (**Figure S2F**) shows approximately equal distribution of male and female samples across racial categories, with the “Other” category containing fewer samples. Because ancestry is a known source of variation in placental biology, we incorporated genomic ancestry principal components (PC1 and PC2 from DNA genotyping) rather than self-reported race categories. This approach directly addresses ancestry-associated bias in gene expression analysis while avoiding socially constructed race categories.

### Variance partition analysis: identification of key covariates

Variance partition analysis quantitatively evaluated the contribution of potential technical and biological covariates to overall gene expression variation, informing DESeq2 model specification for both datasets.

### First trimester: variance components and covariate selection

In first trimester samples (**Figure S3 A and B**), residual (unexplained) variance represented the largest component of gene expression variation, likely reflecting biological heterogeneity between individuals and gene-specific regulatory mechanisms not captured by measured covariates. Of the measured variables, the RNA-seq dataset batch variable explained a modest median variance of 5%, supporting its inclusion in the differential expression model to mitigate confounding by batch effects. Fetal sex explained highly variable amounts of variance across genes, with a long tail of genes in which sex accounted for most of the variance, yet a median variance close to 0%. However, when restricted to autosomes only, sex-related variance decreased dramatically, reaching approximately 25% of the maximum explained variance. This pattern suggests that sex-biased expression in the first trimester placenta is predominantly driven by genes on the sex chromosomes rather than autosomal differences. Fetal sex (male vs female) was retained as the main variable of interest in all models.

Other potential covariates like gestational age (median variance: <2%) and mode of conception (median variance: <1%) explained negligible proportions of expression variance. These variables were therefore excluded from the model to preserve degrees of freedom and avoid unnecessary model complexity. This decision regarding mode of conception is further supported by prior literature ^31^, which found minimal differential gene expression between spontaneously conceived and assisted reproductive technology pregnancies in first trimester chorionic villus, suggesting that mode of conception does not substantially affect the first trimester placental transcriptome.

### Term placenta: variance components and covariate selection

In term placenta samples (**Figure S3 C and D**), residual variance again represented the largest source of expression variation. Nevertheless, the technical contribution was substantially greater than in the first trimester, as the batch variable explained 20–50% of median variance across genes, necessitating its inclusion in the model to prevent systematic bias in differential expression estimates.

Mode of conception was excluded from the term analysis due to a severe sample imbalance (2 of 59 biological samples were conceived through assisted reproduction), which limited statistical power to estimate its effect and would have consumed degrees of freedom without meaningful adjustment.

The lane sequencing variable was omitted from the model because technical replicates were collapsed by summing read counts prior to analysis, thereby rendering lane effects inseparable from biological sample effects. Other biological covariates (parity, pre-pregnancy BMI, gravidity, gestational age, birth weight, maternal age, and method of conception) each explained negligible variance (<5% median) and were therefore excluded from the final model.

Fetal sex explained variable amounts of variance (median 0–25% for most genes), exhibiting a characteristic tail in the distribution indicative of substantial variation across individual genes. As in the first trimester, fetal sex was included in the model as the primary variable of interest.

### PCA: global transcriptional structure

PCA of first trimester and term placenta transcriptomes revealed the dominant sources of variation across different gene selection approaches, complementing the variance partition findings.

### First trimester: batch and sex chromosome effects

In first trimester samples (**Figure S4A-D**), PC1 explained 13–22% of the variance, while PC2 explained 11–18% of the variance. This variance was further increased when restricting the analysis to the 500 most variable genes. Across all analyses, samples showed clear separation by RNA-seq dataset, indicating that batch effects are a major technical driver of global transcriptional structure. Sex-based clustering was prominent in whole-genome analyses (**Figure S4A–B**) but largely disappeared in autosomal-only plots (**Figure S4C–D**), suggesting that sex-specific differences in early placental gene expression are primarily driven by sex chromosome genes.

### Term placenta: batch dominance and attenuated sex clustering

Similarly, term placenta samples (**Figure S4E-H**) showed strong batch effects across all analyses. PC1 explained 26-33% of variance, while PC2 captured 9-19%. Sex-based clustering was most pronounced in the analysis of the top 500 genes (**Figure S4F**) and diminished in autosomal-only plots (**Figure S4G–H**), further indicating the critical role of sex chromosome genes in sex-specific transcriptional differences. PCA did not reveal any obvious clustering by mode of conception, consistent with its negligible variance contribution in the variancePartition analysis. Overall, these findings highlight the importance of considering both sex chromosome contributions and technical batch effects when analysing placental transcriptome data, as sex-based differences may be obscured or amplified depending on whether sex chromosomes are included in the analysis.

### Differential gene expression analysis reveals gestational stage–dependent fetal sex–biased placental expression

Differential gene expression (DGE) analysis results, summarized in **Table 1** and **Figures 2 and 3**, demonstrated sex-biased transcription in both first trimester and term placenta, primarily driven by sex chromosome genes at the single-gene level.

**Figure 2:**
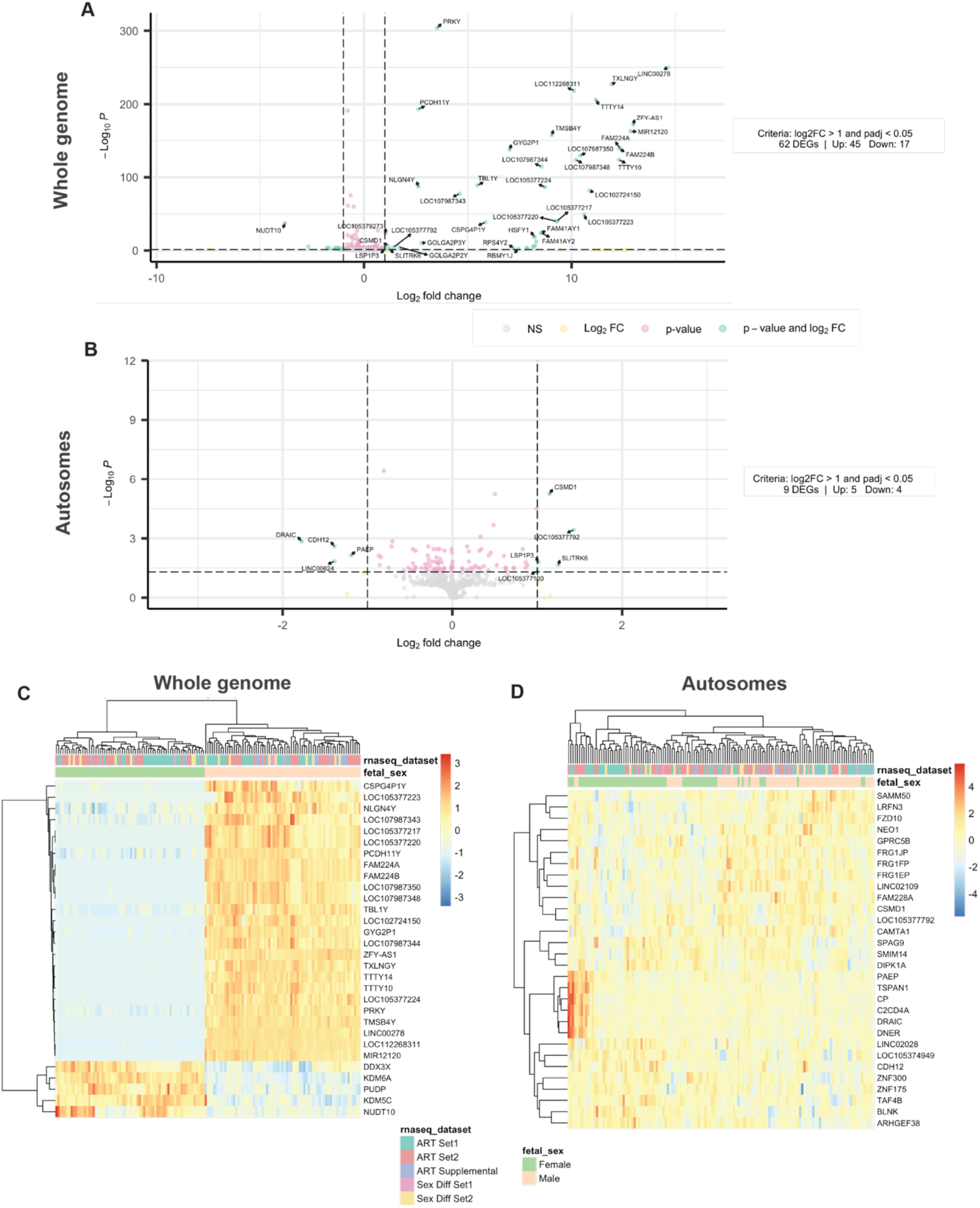
Visualization of differential gene expression for the first trimester database. Panels A and B display volcano plots illustrating differentially expressed genes (DEGs) for the whole genome (A) and autosomes (B). In green genes with both significant adjusted *p-value* (< 0.05) and biologically meaningful effect size (|log2FC| > 1); significant but small effect sizes in pink; large effect size but not statistically significant in yellow; not-significant genes in grey. Panels C and D show heatmaps of the top 30 DEGs for the whole genome (A) and autosomes (B), normalized and scaled across samples. Each row represents an individual gene, and each column represents a placenta sample.

**Figure 3:**
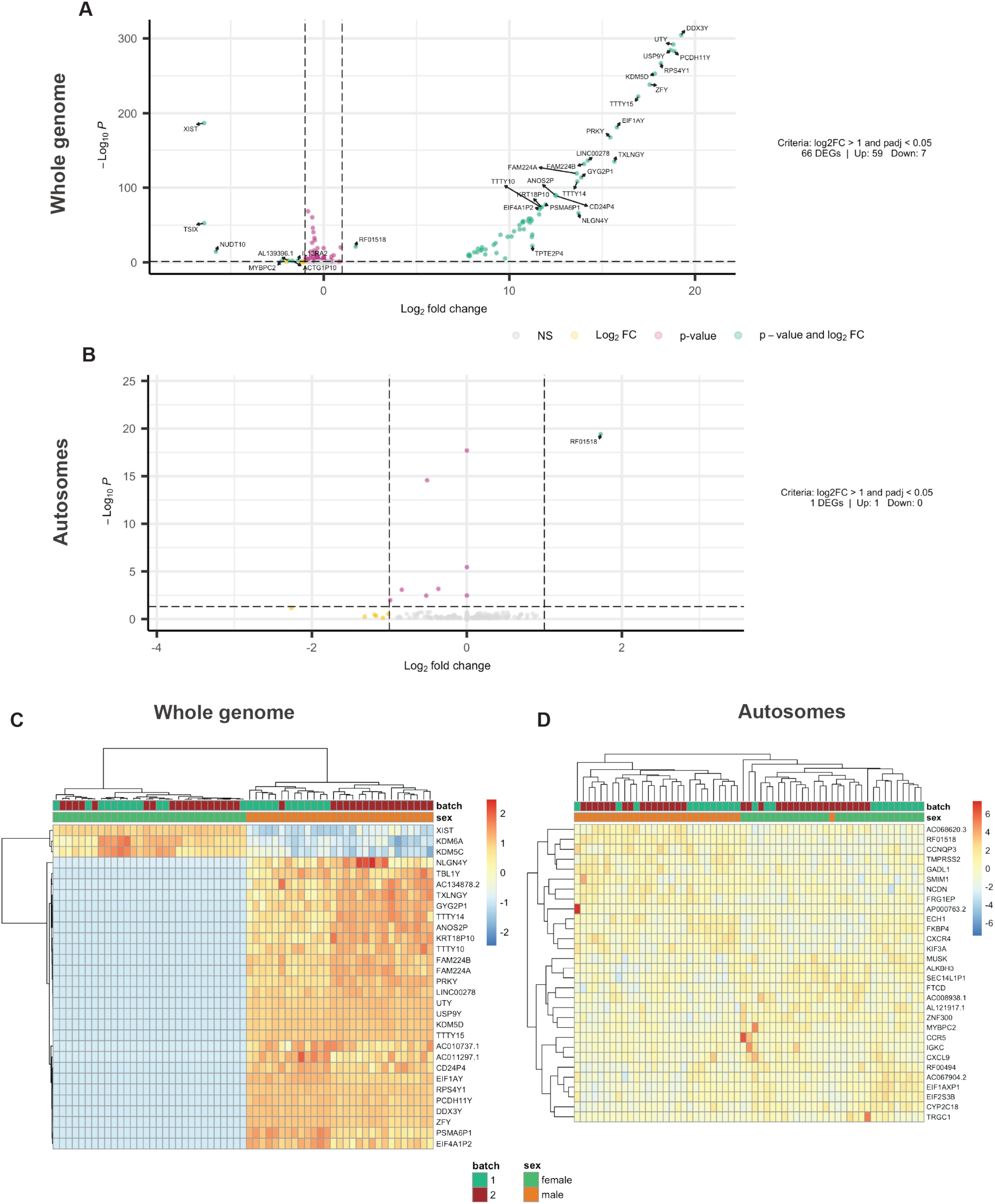
Visualization of differential gene expression for the term database. Panels A and B display volcano plots illustrating differentially expressed genes (DEGs) for the whole genome (A) and autosomes (B). In green genes with both significant adjusted *p-value* (< 0.05) and biologically meaningful effect size (|log2FC| > 1); significant but small effect sizes in pink; large effect size but not statistically significant in yellow; not-significant genes in grey. Panels C and D show heatmaps of the top 30 DEGs for the whole genome (A) and autosomes (B), normalized and scaled across samples. Each row represents an individual gene, and each column represents a placenta sample.

**Table 1:**
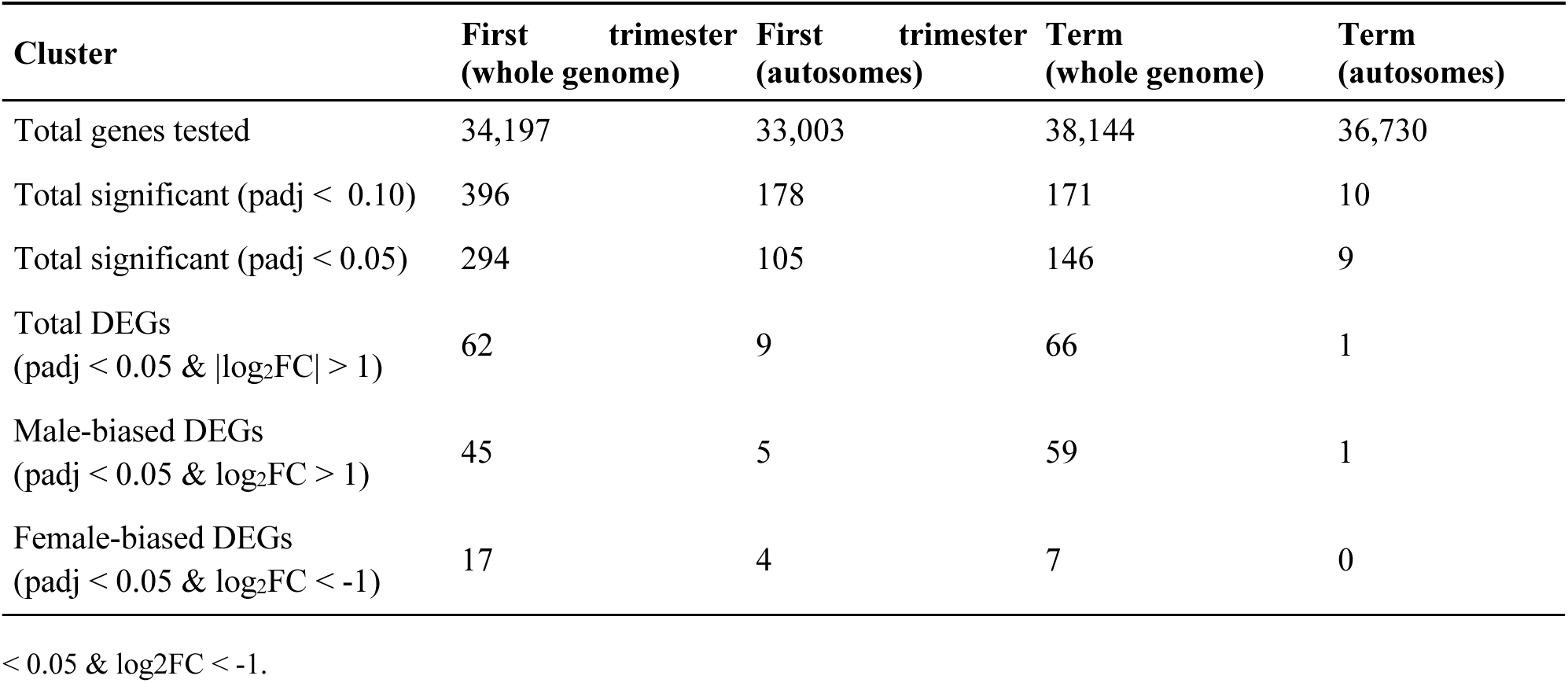
Differentially expressed Genes (DEGs) for the first trimester and term placenta databases. padj < 0.05 considered statistically significant. Male-biased DEGs present padj < 0.05 & log2FC > 1. Female-biased DEGs present padj

In first trimester whole-genome analyses, 62 genes exhibited differential expression between male and female placentas, with 45 upregulated in males and 17 upregulated in females. When restricting to autosomes, only 9 DEGs remained (5 male-biased, 4 female-biased). These patterns are illustrated by volcano plots and heatmaps (**Figure 2**). During the first trimester, sex-biased expression was clearly detectable (**Figure 2A**), and the heatmap revealed a coherent block-wise separation of male and female samples driven by the top 30 DEGs (**Figure 2C**). Upon exclusion of X- and Y-chromosome genes (**Figure 2A and D**), the number of DEGs decreased sharply, and the sex-specific clustering became less distinct, consistent with early sex-biased expression being primarily driven by sex chromosome genes. Consistent with these observations, first trimester whole-genome analyses (**Figure 2A and C**; **Table 1**) indicated that the majority of male-biased DEGs were located on sex chromosomes (40 of 45), while 7 of 17 female-biased DEGs mapped to the X chromosome. Notably, the same five autosomal genes (LOC105377100, SLITRK6, LSP1P3, LOC105377792 and CSMD1) showed male-biased expression in both whole-genome and autosome-only analyses (**Figure 2A and B**), and four autosomal genes (DRAIC, CDH12, PAEP and LINC00624) consistently showed female-biased expression across both analyses. These shared autosomal DEGs likely represent the most robust autosomal contributors to first trimester sex differences.

At term, whole-genome analysis identified 66 DEGs, of which 59 were upregulated in males and 7 in females (**Figure 3A and C**; **Table 1**). Among the male-biased DEGs, only one gene, RF01518, was autosomal; the remaining 58 were Y-linked. Of the seven female-biased DEGs, six mapped to the X chromosome and one, MYBPC2, was autosomal. Considering previous whole-genome analysis results, it is not surprising that the autosomal-only analysis yielded a single DEG, RF01518, upregulated in males (**Figure 3B**), and no obvious sex-specific clustering in the heatmap (**Figure 3D**). Although this gene met the criteria for a true DEG– log2FC > 1 and padj < 0.05–the autosomal-only analysis yielded only a single autosomal DEG, which we judged insufficient for robust DEG-based over-representation or gene-by-gene interpretation. Nevertheless, the full ranked gene list was still used for GSEA. In conjunction with variance-partition and PCA) results (**Figures S3 and S4**), these findings indicate that by late gestation, strong sex-biased transcriptional differences detectable at the single-gene level are predominantly driven by X and Y chromosome genes, whereas autosomal sex differences are more diffuse and generally fall below our conservative DEG threshold.

### Male and female specific hallmarks, pathway and chromosomal enrichment patterns shift from first trimester to term

To explore the functional implications of sex-biased DEG and to assess whether these differences reflect distinct placental developmental or maturation programs, we performed rank-based GSEA.

The rank-based GSEA analysis comparing male versus female first trimester placentas using core pathways and hallmark gene sets (**Figure S5**) revealed distinct pathway enrichment patterns between the fetal sexes. When comparing whole genome (**Figure S5A**) versus autosomal-only analysis (**Figure S5B**), the major patterns remained consistent. However, in the autosomal analysis, we observed slight shifts in the specific pathways while maintaining the overall functional themes. Male fetal sex consistently showed enrichment in epigenetic regulatory pathways (DNA methylation, histone modifications) and gene silencing mechanisms in both analyses. Female fetal sex demonstrated robust enrichment in metabolic processes (amino acid catabolism, fatty acid metabolism) and protein secretion across both analyses, consistent with heightened endocrine and metabolic activity in female placentas. Notably, the *REACTOME_PRE_NOTCH_EXPRESSION_AND_PROCESSING* pathway remained enriched in male placentas in both whole genome and autosomal analyses. In contrast, placental analysis (**Figure S6**) revealed a striking difference from the first trimester results. Unlike the first trimester patterns, where male placentas showed enrichment in epigenetic regulation, term placental transcriptomes exhibited predominantly female-enriched pathways, with no significantly enriched pathways in male placentas among the top 20 results. Female term placentas exhibited robust enrichment across multiple immune and inflammatory pathways, with NES consistently around −2. Excluding sex chromosomes from the analysis (autosomes only) identified only one DEG was identified (padj < 0.05, |log2FC| > 1), and autosomal-only GSEA did not yield any significantly enriched pathways at FDR < 0.05 (data not shown). Therefore, we focused on whole-genome pathway results for the term dataset, which capture the dominant sex chromosome-driven signal while still incorporating autosomal contributions.

Our chromosome-level GSEA analysis of first trimester placentas revealed distinct patterns of enrichment between whole genome (**Figure S7A**) and autosomal-only analyses (**Figure S7B**) as expected. In the whole genome analysis (**Figure S7A**), Y chromosome regions showed the strongest male-biased enrichment, with chrYq11 and chrYp11 having the highest normalized enrichment scores (NES > 2.5). Several autosomal regions also showed significant male-biased enrichment, notably chr6p22, with an NES approaching 2.0. Female-biased enrichment in the whole genome analysis was primarily observed on the X chromosome (chrXq13, chrXp11, and chrXp22). When restricting the analysis to autosomes only (Figure S7B), a substantially different pattern emerged. The most strongly male-enriched autosomal regions were chr6p22, chr22q13, and chr9q34, with NES values approximately 2.0. Female-enriched autosomal regions included chr12q22, chr6q14, chr1p31, and chr5q13, with NES values lower than −2.0. These positional enrichments likely reflect clusters of sex-biased genes within these regions; however, they do not identify specific regulatory elements. For term placentas, the whole genome analysis (Figure S8) again revealed that Y chromosome regions exhibited the strongest male-biased enrichment, with chrYq11 and chrYp11 demonstrating the highest normalized enrichment scores (NES > 3.0). Female-biased enrichment maintained strong signals on the X chromosome (chrXq13, chrXp11, chrXp22). Autosomal regions also showed female bias, including chr4q25 and chr14q22, however, with NES values slightly lower than −2.

### Gestational reversal of sex differences in placental vascular and extracellular matrix developmental signalling

Considering the hypothesis of sex differences in placental vascularization and maturation, gene sets enriched for vascular and barrier-related terms were searched.

In the first trimester whole-genome analysis, five significantly enriched vascular and barrier-related pathways were identified (**Figure 4A**), all exhibiting higher activity in male placentas (**Table 2**). The strongest signal corresponded to “Reactome Pre-NOTCH Expression and Processing” (NES > 2.5, FDR < 0.01). When the analysis was restricted to autosomes, this signal intensified rather than diminished, with more than twenty vascular- and barrier-related pathways showing significant male-enriched activity (**Figure 4B**; **Table 3**). Despite the removal of all X- and Y-linked genes, the autosome-only leading-edge subsets remained highly coherent and mapped into a limited number of recurring pathway modules. Multiple NOTCH-related pathways, including NOTCH signalling, NOTCH-3 intracellular domain regulation, and NOTCH-overexpression modules, were consistently enriched. In parallel, integrin-extracellular matrix (ECM) and adhesion pathways (αVβ3 and α9β1 integrin modules, platelet adhesion to collagen, and non-integrin ECM interactions) were prominently represented, driven by collagens (COL1A1/A2, COL3A1, COL4A1/A2, COL5A1/A2/A3, COL6A1/A2/A3), integrins (ITGA1/2/4/2B), adhesion molecules (ICAM1/2/3, PECAM1), and angiogenic mediators (VTN, KDR, THBS1). Additionally, several RUNX1-associated megakaryocyte and platelet-involved pathways were enriched in males. Finally, chromatin-associated programs, including RNA Polymerase I and III transcription and promoter-escape modules, remained strongly represented, with leading-edge genes dominated by canonical histone clusters (H2A, H2B, H3, and H4 families), reflecting the structure of these gene sets.

**Figure 4:**
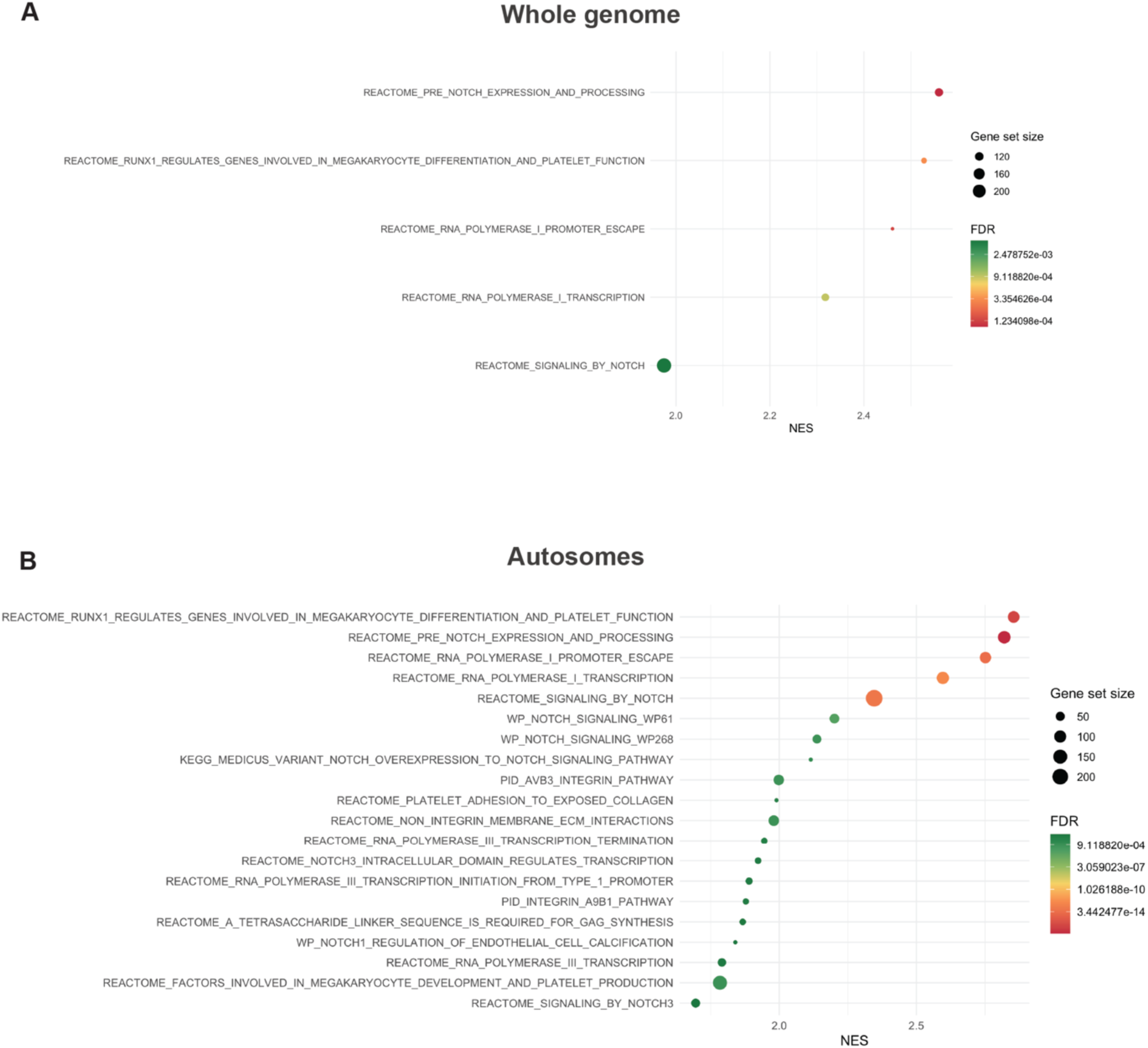
Rank-based gene set enrichment analysis (fgsea) of fetal sex-biased gene expression in first trimester placenta focusing on vascular development and barrier function pathways. (A) Whole genome and (B) Autosomes. Positive NES indicates pathways upregulated in males, while negative NES indicates pathways upregulated in females. Point size = gene set size; color = FDR. MSigDB gene sets (15–500 genes); FDR < 0.05 was considered statistically significant. Pathway databases queried: Hallmark (H), KEGG (C2:CP:KEGG_MEDICUS), Reactome (C2:CP:REACTOME), WikiPathways (C2:CP:WIKIPATHWAYS), BioCarta (C2:CP:BIOCARTA), and Pathway Interaction Database (C2:CP:PID) from the Molecular Signatures Database (MSigDB v25.1.1).

**Table 2:**
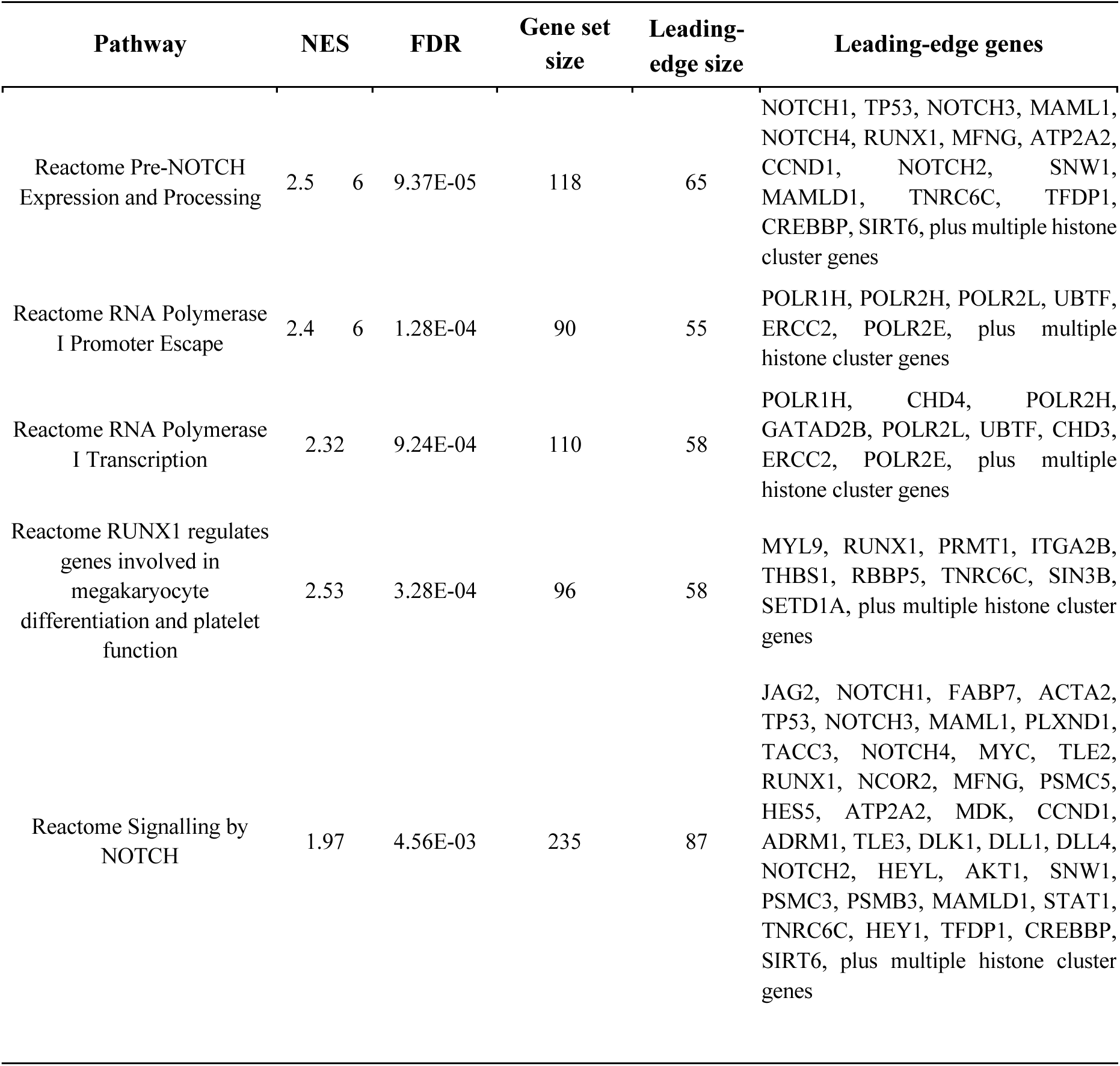
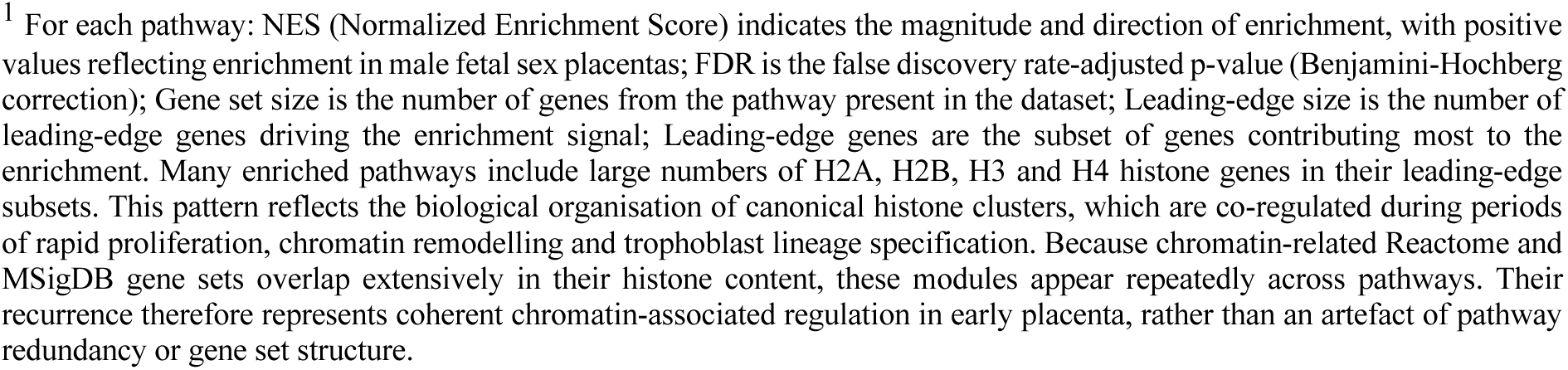
List of vascular-focused enriched pathways and leading-edge genes identified from the whole genome of first trimester placenta.^1^.

| Pathway | NES |  | FDR | Gene set size | Leading-edge size | Leading-edge genes |
| --- | --- | --- | --- | --- | --- | --- |
| Reactome Pre-NOTCH Expression and Processing | 2.5 | 6 | 9.37E-05 | 118 | 65 | NOTCH1, TP53, NOTCH3, MAML1, NOTCH4, RUNX1, MFNG, ATP2A2, CCND1, NOTCH2, SNW1, MAMLD1, TNRC6C, TFDP1, CREBBP, SIRT6, plus multiple histone cluster genes |
| Reactome RNA Polymerase I Promoter Escape | 2.4 | 6 | 1.28E-04 | 90 | 55 | POLR1H, POLR2H, POLR2L, UBTF, ERCC2, POLR2E, plus multiple histone cluster genes |
| Reactome RNA Polymerase I Transcription | 2.32 |  | 9.24E-04 | 110 | 58 | POLR1H, CHD4, POLR2H, GATAD2B, POLR2L, UBTF, CHD3, ERCC2, POLR2E, plus multiple histone cluster genes |
| Reactome RUNX1 regulates genes involved in megakaryocyte differentiation and platelet function | 2.53 |  | 3.28E-04 | 96 | 58 | MYL9, RUNX1, PRMT1, ITGA2B, THBS1, RBBP5, TNRC6C, SIN3B, SETD1A, plus multiple histone cluster genes |
| Reactome Signalling by NOTCH | 1.97 |  | 4.56E-03 | 235 | 87 | JAG2, NOTCH1, FABP7, ACTA2, TP53, NOTCH3, MAML1, PLXND1, TACC3, NOTCH4, MYC, TLE2, RUNX1, NCOR2, MFNG, PSMC5, HES5, ATP2A2, MDK, CCND1, ADRM1, TLE3, DLK1, DLL1, DLL4, NOTCH2, HEYL, AKT1, SNW1, PSMC3, PSMB3, MAMLD1, STAT1, TNRC6C, HEY1, TFDP1, CREBBP, SIRT6, plus multiple histone cluster genes |

<sup>1</sup> For each pathway: NES (Normalized Enrichment Score) indicates the magnitude and direction of enrichment, with positive values reflecting enrichment in male fetal sex placentas; FDR is the false discovery rate-adjusted p-value (Benjamini-Hochberg correction); Gene set size is the number of genes from the pathway present in the dataset; Leading-edge size is the number of leading-edge genes driving the enrichment signal; Leading-edge genes are the subset of genes contributing most to the enrichment. Many enriched pathways include large numbers of H2A, H2B, H3 and H4 histone genes in their leading-edge subsets. This pattern reflects the biological organisation of canonical histone clusters, which are co-regulated during periods of rapid proliferation, chromatin remodelling and trophoblast lineage specification. Because chromatin-related Reactome and MSigDB gene sets overlap extensively in their histone content, these modules appear repeatedly across pathways. Their recurrence therefore represents coherent chromatin-associated regulation in early placenta, rather than an artefact of pathway redundancy or gene set structure.

**Table 3:**
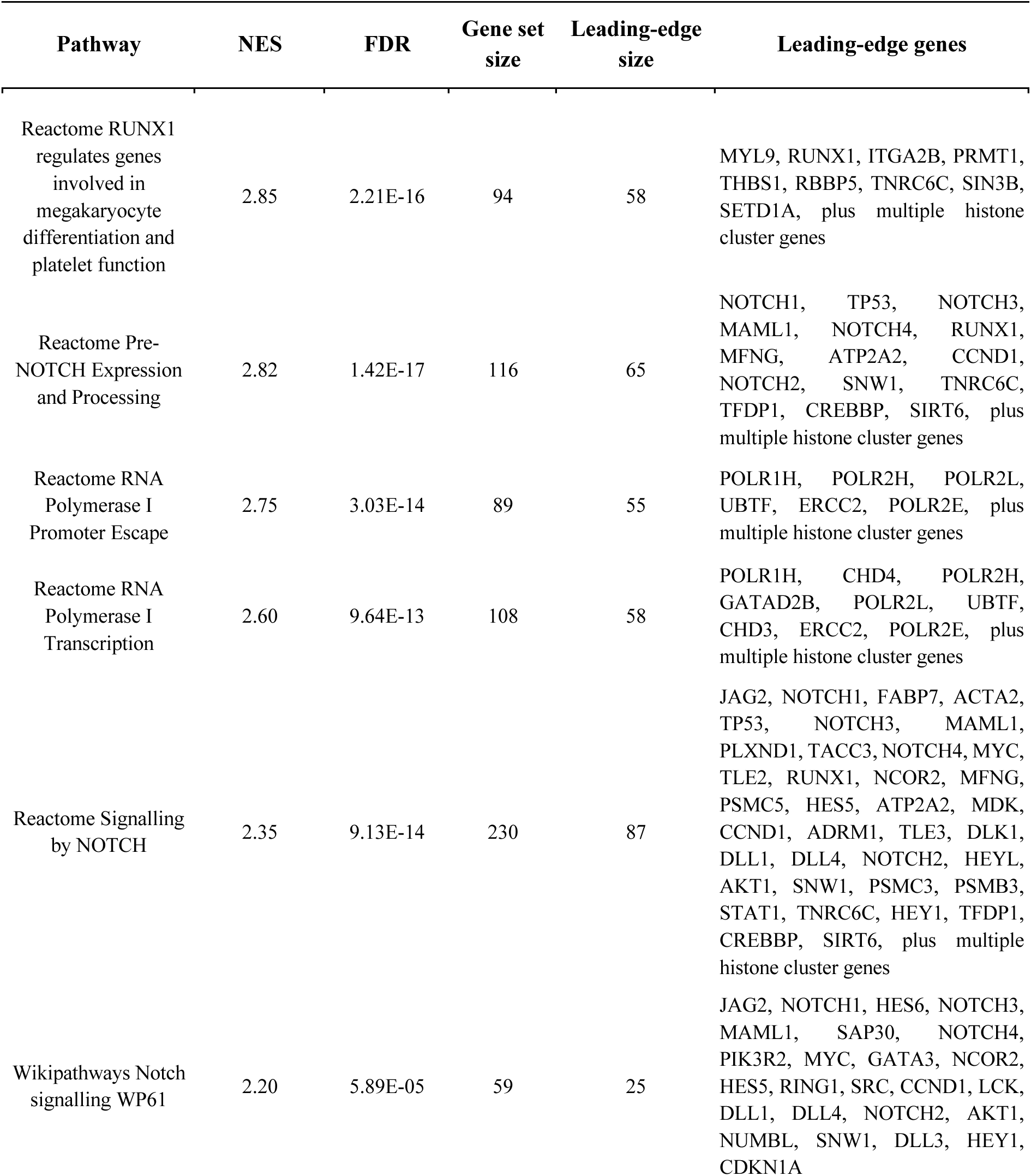
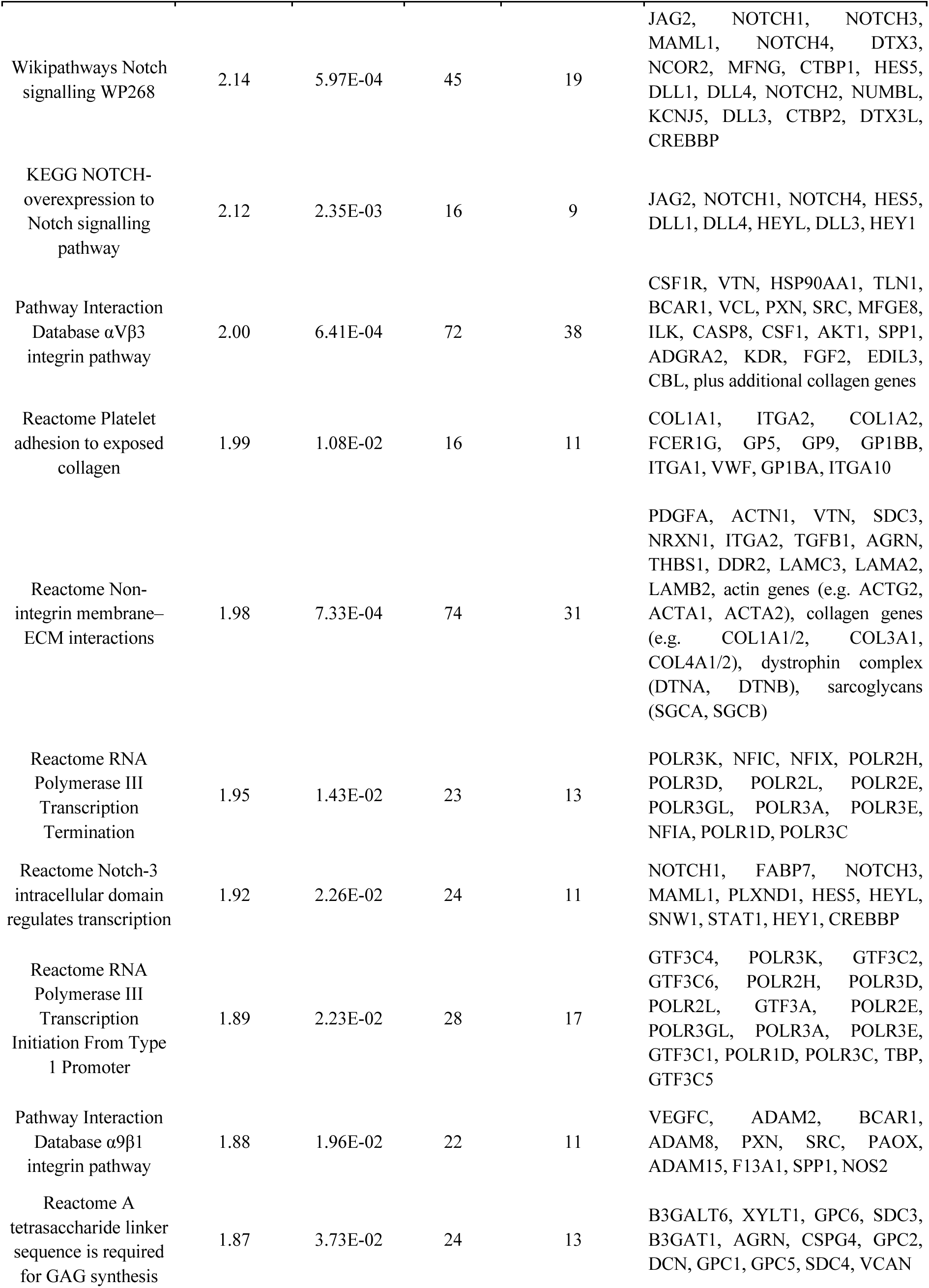
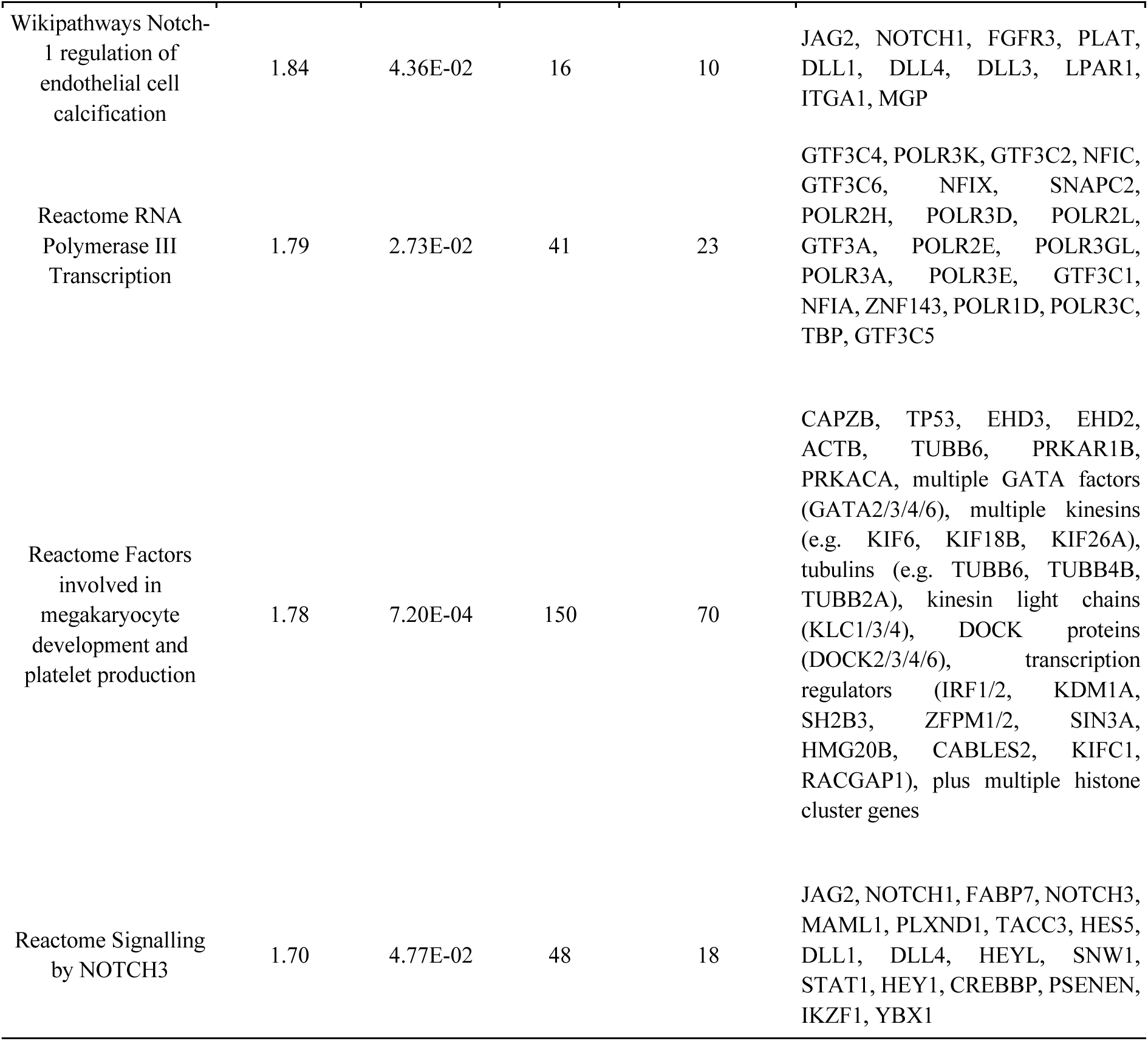
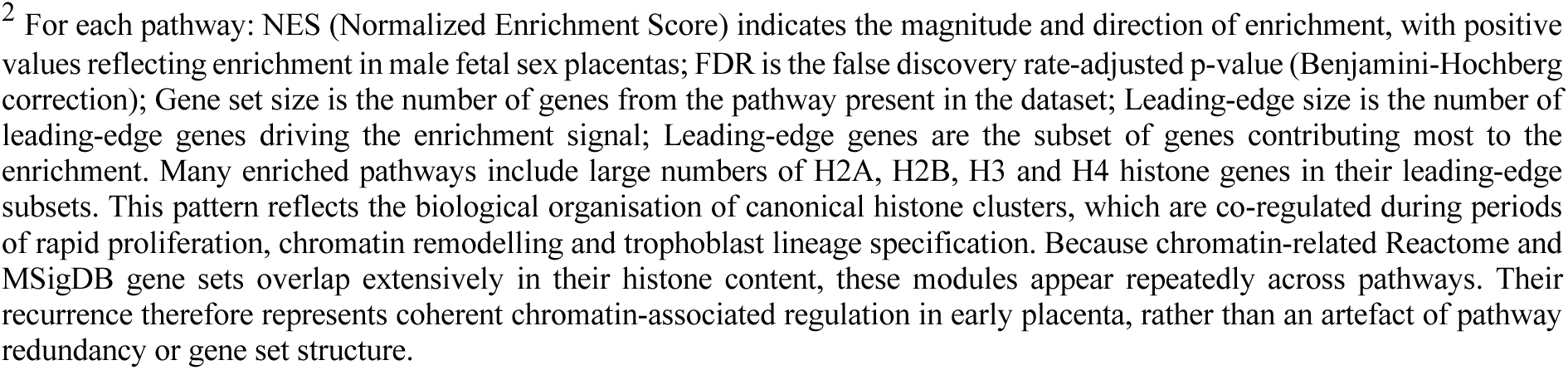
List of vascular-focused enriched pathways and leading-edge genes identified from first trimester placental data (autosomes). The table displays the top 20 vascular and barrier pathways exhibiting the largest absolute normalized enrichment scores (NES), with a Hallmark/curated pathways with FDR < 0.05.^2^

At term, whole-genome analysis identified three vascular- and barrier-related pathways that reached statistical significance (**Figure 5**; **Table 4**); however, the direction of enrichment was reversed compared to the first trimester. These pathways showed female-enriched activity, encompassing modules related to vascular wall interactions and inflammatory or anti-inflammatory responses. Gene-set sizes were intermediate (approximately 100–190 genes), and leading-edge subsets were relatively broad (around 50–90), indicating that these vascular and immune signals reflect coordinated regulation of immunoglobulins, Fc receptors, integrins and leukocyte adhesion molecules rather than highly restricted focal gene effects.

**Figure 5:**
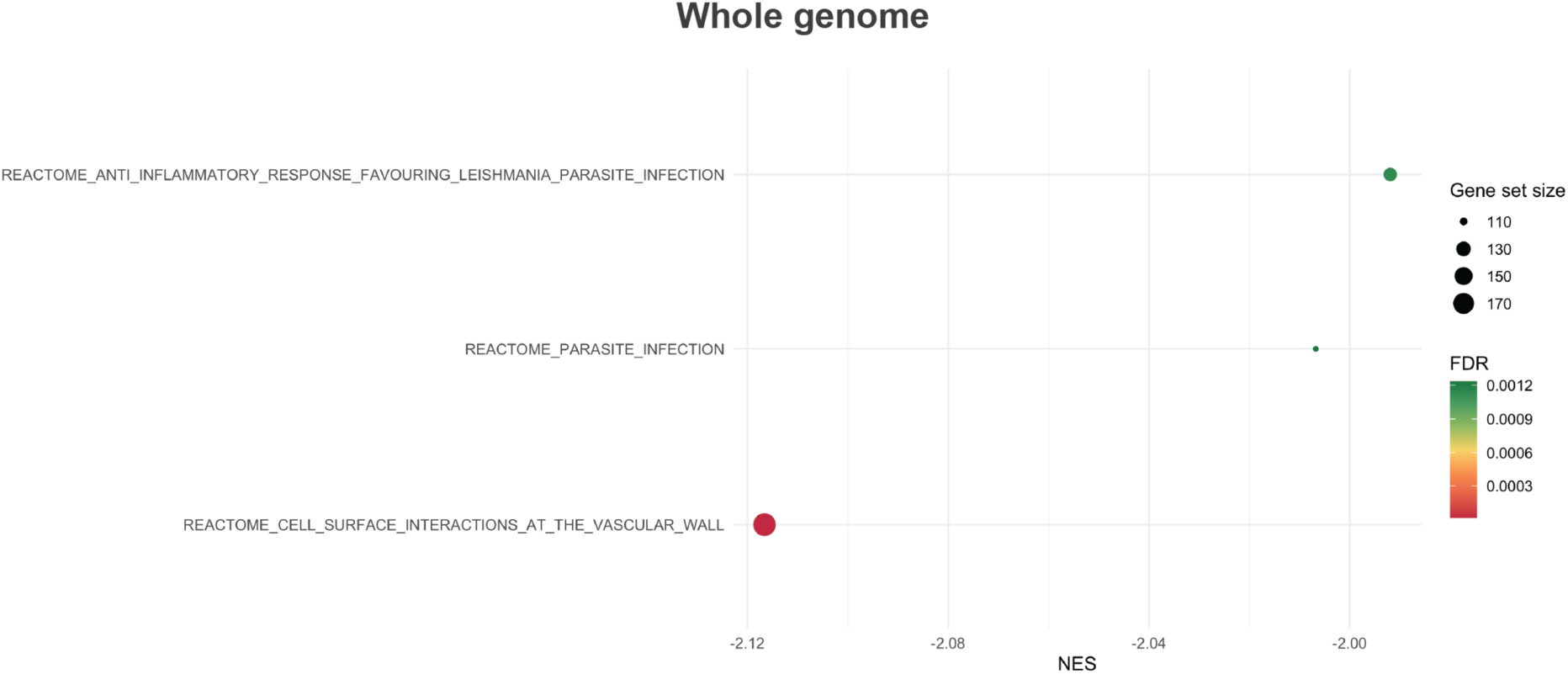
Rank-based gene set enrichment analysis (fgsea) of fetal sex-biased gene expression in term placenta (whole genome) focusing on vascular development and barrier function pathways. Positive NES = pathways upregulated in males; negative NES = pathways upregulated in females. Point size = gene set size; color = FDR. MSigDB gene sets (15–500 genes); FDR threshold of < 0.05 was considered statistically significant. Pathway databases queried: Hallmark (H), KEGG (C2:CP:KEGG_MEDICUS), Reactome (C2:CP:REACTOME), WikiPathways (C2:CP:WIKIPATHWAYS), BioCarta (C2:CP:BIOCARTA), and Pathway Interaction Database (C2:CP:PID) from the Molecular Signatures Database (MSigDB v25.1.1).

**Table 4:**
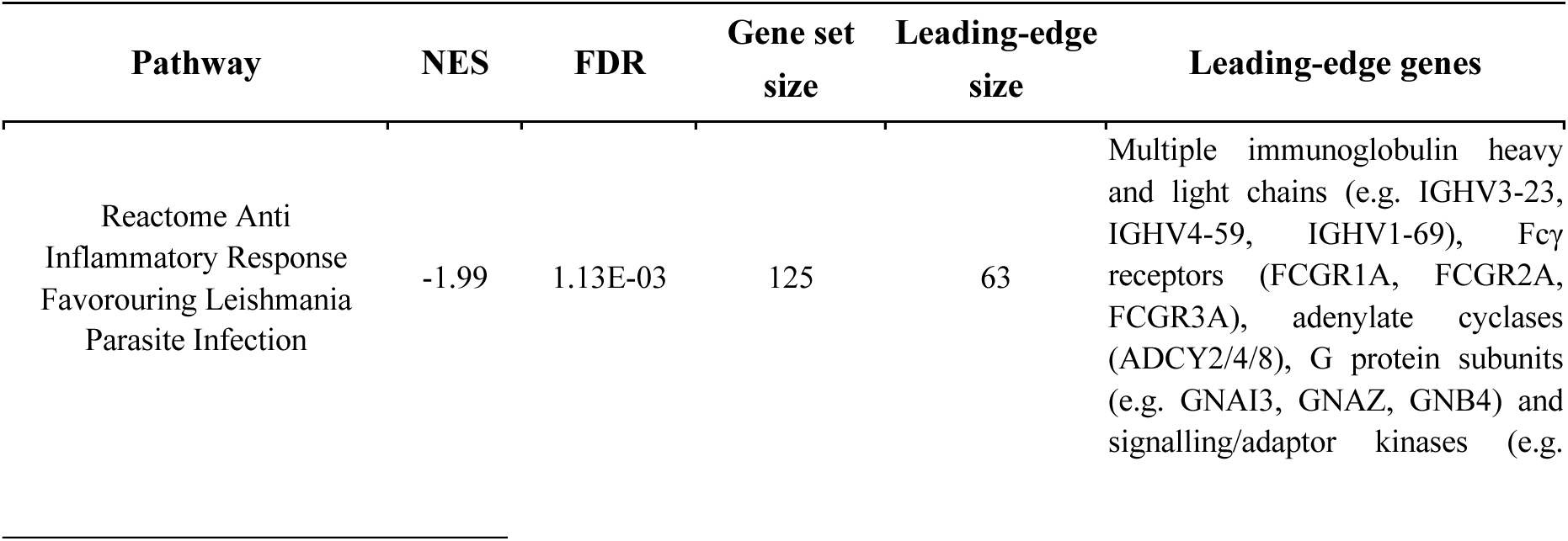
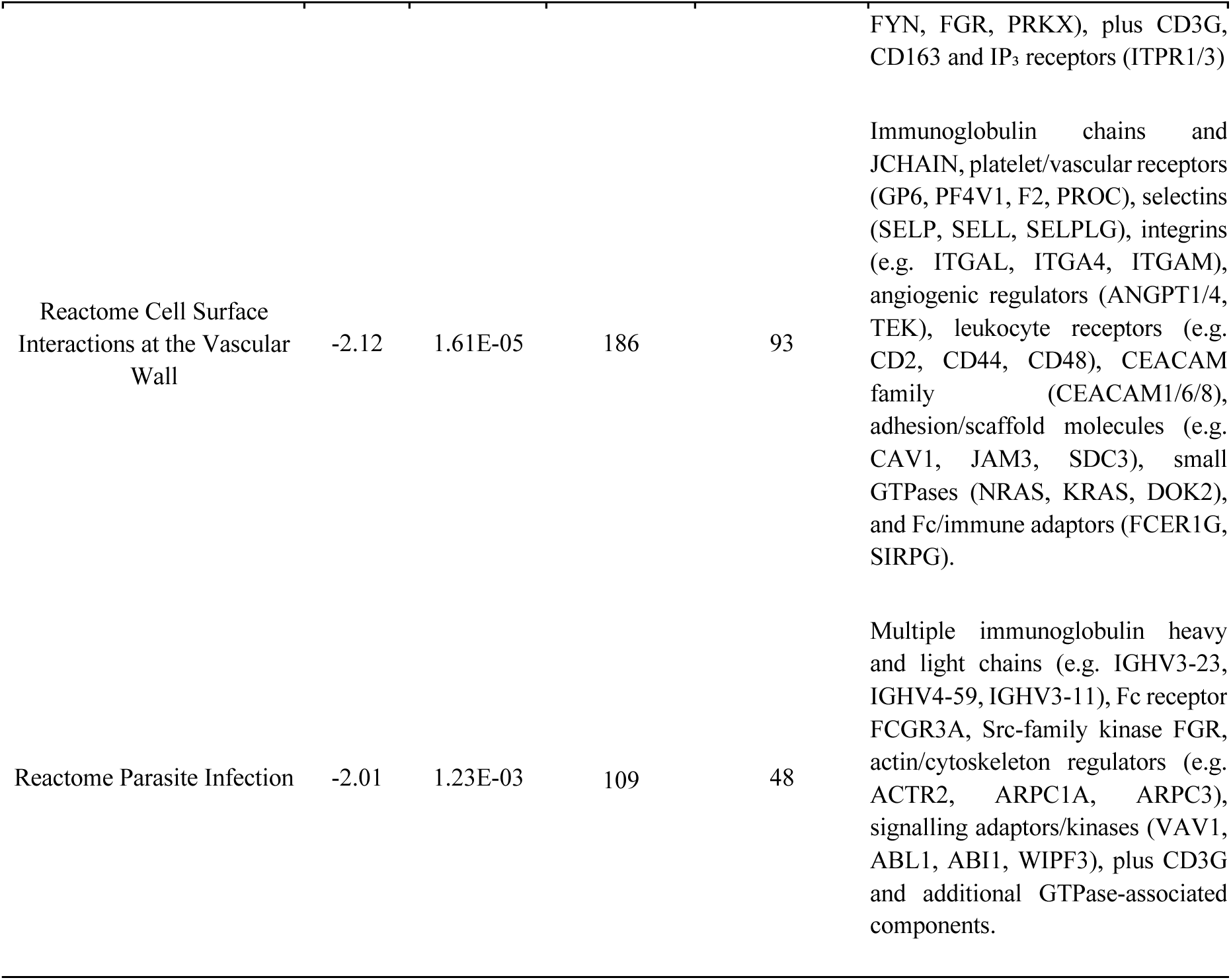
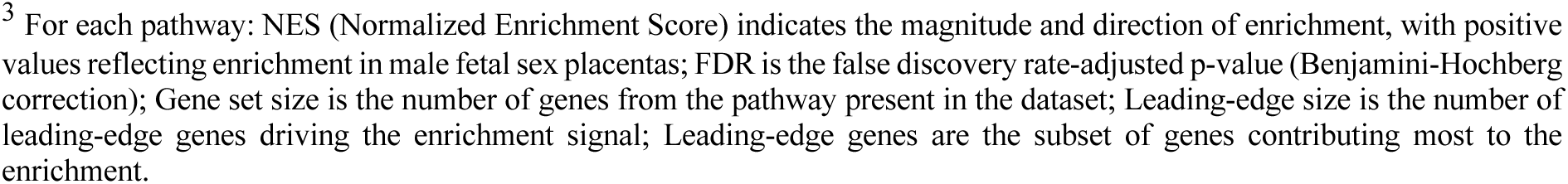
List of vascular-focused enriched pathways and leading-edge genes identified from the whole genome of term placenta.^3^.

| Pathway | NES | FDR | Gene set size | Leading-edge size | Leading-edge genes |
| --- | --- | --- | --- | --- | --- |
| Reactome Anti Inflammatory Response Favouring Leishmania Parasite Infection | -1.99 | 1.13E-03 | 125 | 63 | Multiple immunoglobulin heavy and light chains (e.g. IGHV3-23, IGHV4-59, IGHV1-69), Fcγ receptors (FCGR1A, FCGR2A, FCGR3A), adenylate cyclases (ADCY2/4/8), G protein subunits (e.g. GNAI3, GNAZ, GNB4) and signalling/adaptor kinases (e.g. |

<sup>3</sup> For each pathway: NES (Normalized Enrichment Score) indicates the magnitude and direction of enrichment, with positive values reflecting enrichment in male fetal sex placentas; FDR is the false discovery rate-adjusted p-value (Benjamini-Hochberg correction); Gene set size is the number of genes from the pathway present in the dataset; Leading-edge size is the number of leading-edge genes driving the enrichment signal; Leading-edge genes are the subset of genes contributing most to the enrichment.

| Pathway | NES | FDR | Gene set size | Leading-edge size | Leading-edge genes |
| --- | --- | --- | --- | --- | --- |
|  |  |  |  |  | FYN, FGR, PRKX), plus CD3G, CD163 and IP<sub>3</sub> receptors (ITPR1/3) |
| Reactome Cell Surface Interactions at the Vascular Wall | -2.12 | 1.61E-05 | 186 | 93 | Immunoglobulin chains and JCHAIN, platelet/vascular receptors (GP6, PF4V1, F2, PROC), selectins (SELP, SELL, SELPLG), integrins (e.g. ITGAL, ITGA4, ITGAM), angiogenic regulators (ANGPT1/4, TEK), leukocyte receptors (e.g. CD2, CD44, CD48), CEACAM family (CEACAM1/6/8), adhesion/scaffold molecules (e.g. CAV1, JAM3, SDC3), small GTPases (NRAS, KRAS, DOK2), and Fc/immune adaptors (FCER1G, SIRPG). |
| Reactome Parasite Infection | -2.01 | 1.23E-03 | 109 | 48 | Multiple immunoglobulin heavy and light chains (e.g. IGHV3-23, IGHV4-59, IGHV3-11), Fc receptor FCGR3A, Src-family kinase FGR, actin/cytoskeleton regulators (e.g. ACTR2, ARPC1A, ARPC3), signalling adaptors/kinases (VAV1, ABL1, ABI1, WIPF3), plus CD3G and additional GTPase-associated components. |

This demonstrates a dynamic, gestation-dependent shift in sex-biased placental developmental programs, with male placentas exhibiting early enrichment of vascular expansion pathways (NOTCH signaling, angiogenesis, integrin-ECM interactions, and extracellular matrix remodeling), while female placentas show predominant enrichment of vascular maintenance, barrier function, and immune-vascular integration pathways at term.

### Sex-biased transcriptional programs highlight convergent patterns across multiple analytical scales

Across all analyses, several consistent patterns emerged. First, sex-biased transcription was detected at both gestational stages, but was predominantly driven by X- and Y-linked genes at the single-gene level, with very few autosomal DEGs, particularly at term. Second, variance partitioning and PCA showed that batch was a major technical source of variation in both datasets, and that sex-related variance and sex-based clustering were stronger in first trimester placentas and largely dependent on the inclusion of sex chromosome genes. Third, pathway-level analyses of ranked gene lists from first trimester placentas revealed male-enriched epigenetic, chromatin, and vascular/NOTCH-related pathways, and female-enriched metabolic and secretion pathways, with broadly similar themes in both whole-genome and autosome-only analyses. Fourth, in term placentas, pathways and hallmark GSEA identified predominantly female-enriched immune and inflammatory pathways, with no significantly enriched pathways in males among the top-ranked results. Finally, chromosome-level and vascular-focused GSEA demonstrated that sex-biased enrichment of Y and X chromosomal regions was accompanied by specific autosomal positional and vascular pathway enrichments at both stages, despite the paucity of autosomal DEGs, providing a coherent transcriptomic framework for the developmental and vascular sex differences addressed in the Discussion section.

## Discussion

Despite the central role of this vascular tree in shaping placental growth and pregnancy outcome, and the evidence that fetal sex imprints placental structure, function, and molecular signalling, how sex remodels the placental vasculature across gestation remains largely uncharted. This gap in knowledge represents a critical blind spot in developmental and reproductive biology. To address this gap, we analysed RNA-Seq data from two independent cohorts spanning early and late gestation, utilizing harmonized and single-gene differential expression analysis alongside pathways enrichment strategies to determine how sex-biased transcriptional programs change over time, with a particular focus on vascular and barrier-related pathways.

Across both developmental windows, sex-biased transcription was readily detectable, but its architecture differed markedly by gestational stage. In the first trimester, placentas from male fetuses showed pronounced enrichment of epigenetic regulatory pathways, including gene sets linked to DNA methylation and chromatin regulation (e.g., chromatin remodelers and transcriptional regulators such as CHD3, CHD4, GATAD2B, SETD1A, SIN3B, PRMT1, and CREBBP), as well as multiple canonical histone cluster genes contributing to RNA polymerase and chromatin-associated modules. Although our analysis is transcriptional and does not directly assess DNA methylation or histone modifications, prior placental literature has implicated sex-linked regulation of imprinted loci (e.g., SNRPN, PEG10, MEST) and broader chromatin-state modulation, including repressive histone marks such as H3K27me3 and H3K9me3 ^39^, situating our chromatin-associated transcriptional findings within a broader epigenetic framework. Meanwhile, female placentas demonstrated enrichment of metabolic pathways, particularly fatty acid metabolism (e.g., β-oxidation via ACADL), amino acid catabolism, and protein secretion/ER processing. These chromatin and transcription signatures are consistent with the heightened transcriptional plasticity characteristic of early placental development, as reported in transcriptomic studies showing sex-specific regulation of chromatin genes. At term, unlike the first trimester, where both sexes showed robust pathway enrichment, term placentas exhibited predominantly female-enriched immune and inflammatory pathways, including interferon signalling, B-cell receptor activation, and immunoregulatory interactions, with no male-enriched pathways among the top-ranked global results and minimal autosomal sex-differential gene expression at the single-gene level. Our findings suggest that placental sexual dimorphism is developmentally dynamic, with sex-specific molecular programs at early and late pregnancy that may be influencing overall fetal development. We hypothesise that male placentas may adopt a developmental trajectory that prioritizes rapid somatic growth, potentially allocating proportionally fewer transcriptional resources to antiviral and immunoregulatory defenses. Such a configuration could limit flexibility in responding to maternal inflammatory or infectious challenges. Consistent with this model, epidemiological and biometric data show that male fetuses are, on average, larger than females throughout gestation, exhibiting greater crown–rump length, head and abdominal circumferences, higher birthweight, and an elevated fetal-to-placental weight ratio ^38^. Within this framework, maternal bacterial infections during pregnancy are associated with increased risk of psychosis in offspring, with males showing nearly three-fold higher odds than females, indicating a sex-dependent sensitivity to prenatal immune challenge. Compounding this, male fetuses also appear more prone to downstream neurodevelopmental vulnerability to early-life infection burden and later psychopathology ^40^. On the other hand, female placentas appear to integrate metabolic regulation with immune preparedness across gestation, thereby buffering fetal development under stress and contributing to a more resilient placental phenotype. In human studies of spontaneous preterm birth, female placental tissues maintain immunometabolic profiles biased toward anti-inflammatory states and exhibit comparatively restrained transcriptomic responses to pathological stressors. Male placentas, by contrast, show widespread gene expression alterations in pathways related to energy metabolism and inflammation and exhibit greater perturbation under pathophysiological stress ^41^.

A key biological insight from variance partition analysis of first trimester placentas is that sex chromosomes are the primary drivers of sex-associated transcriptional variation. When sex-chromosome features are excluded, the variance attributable to fetal sex among sex-biased genes is greatly reduced. For many sex-linked transcripts, fetal sex accounts for approximately 50–100% of expression variance in the genome-wide model, but this drops to less than 25% when sex chromosomes are removed. At term, exclusion of sex chromosomes substantially reduced detectable sex-biased signal below a conservative threshold; however, and the overall magnitude of sex-associated variance, attributable to fetal sex, was lower than in early gestation. Physiologically, this pattern is consistent with an early gestational window in which placental sex differences are driven primarily by sex-chromosome dosage, underscoring the central role of X/Y complement in early development ^42–44^ and then becomes progressively attenuated across gestation. In early gestation, sex-associated variance is plausibly dominated by sex-chromosome dosage effects, including male-specific Y expression and female-biased expression of X-linked genes that escape X-chromosome inactivation ^45,46^. By term, the smaller fraction of variance attributable to fetal sex, together with the persistence of modest autosomal sex-associated variance after excluding sex chromosomes, points to a relative enrichment for autosomal encoded regulatory differences. Such later differences may arise from developmental remodelling of placental cell states and from sex-biased endocrine, immune, and vascular signalling programs implicated in sex-specific fetal growth trajectories and susceptibility to pregnancy complications ^47,48^.

To then interrogate how sex-specific transcriptional differences extend to placental vascular biology, we performed functional enrichment analysis focused on vascular and barrier-related pathways. This analysis revealed a coherent picture in which male and female placentas adopt distinct vascular strategies that are both temporally and functionally tuned. In early gestation, there is strong male-enriched activation of NOTCH-related pathways. Consistent with our results, Cvitic and colleagues ^49^ reported male-biased placental gene expression and identified Notch signalling as a pathway influenced by fetal sex. The Notch signalling pathway regulates cell proliferation and death ^50^, endothelial tip–stalk specification, arterial identity, and labyrinthine vascular formation during placentation ^51,52^. Perturbation of endothelial Jagged/Delta–Notch interactions disrupt decidual angiogenesis and spiral-artery remodelling ^53^. A placenta characterised by strong NOTCH–integrin–RUNX1-associated signalling may be particularly susceptible to perturbations such as hypoxia, inflammation, or altered maternal vascular resistance, because disruption at any node can reverberate through interconnected processes, including endothelial differentiation, matrix remodelling, and platelet-endothelial crosstalk. Indeed, impaired NOTCH signalling ^54^ and reduced JAG1 expression are observed in preeclamptic pregnancies, a condition strongly associated with adverse neurodevelopmental outcomes ^55^. The parallel enrichment of integrin-ECM, adhesion, and platelet/megakaryocyte programs supports the view that the male placental vasculature is configured as an actively remodelling and angiogenic niche, where endothelial cells, matrix components, and platelets cooperate to rapid vessel branching, stabilize nascent vessels, and regulate barrier properties, favouring vascular expansion rather than simply increasing trophoblast mass. Human fetoplacental endothelial cells cultured on placental villous stromal cell–derived matrices exhibit profound changes in migration and proliferation, and FGR-derived matrices impair angiogenic behaviour in both control and FGR endothelial cells, underscoring that villous stromal ECM architecture and integrin signalling are primary determinants of ongoing fetoplacental angiogenesis (Ji et al., 2021). Collagen and laminin-rich basement membranes provide the principal structural scaffold for fetoplacental capillaries and villous surfaces. Experimental blockade of collagen at the fetal-maternal interface impairs extravillous trophoblast invasion in vitro, illustrating how basement-membrane composition integrates with trophoblast and endothelial biology within the functional placental barrier ^56^. This is in line with data showing that villous stromal ECM architecture and integrin signalling are key determinants of fetoplacental angiogenesis and are disrupted in severe FGR ^11^, suggesting that failure of early placental vascular adaptation may have more pronounced downstream consequences for fetoplacental perfusion and fetal brain development in males.

By term, this landscape shifts: female placentas demonstrate relatively higher activity in vascular wall interactions and inflammatory or anti-inflammatory modules, whereas no vascular pathways remain male-enriched among the statistically robust signals. This pattern suggests a transition from broad, growth-oriented angiogenic programs toward more focused maintenance- and defence-oriented functions in females. Protein-level analyses similarly document sex differences in placental cytokines ^57^, angiogenic factors ^54^ and glucocorticoid-related proteins ^58^, supporting the concept that females engage more flexible vascular-immune adaptation as gestation progresses.

Longitudinal profiling of maternal plasma across gestation shows that women carrying a male fetus exhibit higher circulating levels of pro-angiogenic factors and a more pro-inflammatory cytokine milieu, whereas pregnancies with female fetuses are associated with relatively higher levels of regulatory cytokines and more complex patterns of angiogenesis-related mediators ^59^. Preferential investment in vascular expansion and endothelial signalling during early gestation is consistent with a placental strategy oriented toward supporting accelerated fetal growth. Such a configuration likely optimizes oxygen and nutrient transfer under baseline conditions, but it may also limit the capacity to redirect resources toward immune or stress-adaptive pathways when the intrauterine environment becomes adverse. In this context, our findings point to a stage-dependent and functionally divergent pattern of sex-specific transcriptional programming within placental vascular biology. Whereas term male placentas show relatively muted enrichment of vascular and immune-remodelling pathways, females display selective upregulation of barrier and immune-vascular integration programs, reflecting a balance between growth prioritization and immunological vigilance.

It is important to note that our analysis of the term placenta dataset builds directly upon the work of Olney et al ^26^, who first characterised sex-biased expression patterns in this cohort and reported a robust female-enriched immune signatures at term. Our findings are broadly consistent with these core observations; however, we also report some differences, including a relatively stronger male-biased component in our whole-genome DESeq2 models, which likely reflect analytical focus and modelling assumptions rather than conflicting biology. Olney et al. modelled birthweight and sequencing lane, applied FPKM filtering, and used limma-voom to derive DEG calls, whereas our framework used raw counts with DESeq2, collapsed technical replicates such that lane attribution was resolved at the sample level, excluded birthweight as a sex-influenced downstream phenotype, and complemented DEG-level analysis with rank-based pathway enrichment. Birthweight was excluded because fetal sex is an upstream determinant of placental function and fetal growth, so adjusting for birthweight may confound some biologically meaningful sex-linked expression differences, particularly those related to growth trajectories. These methodological distinctions emphasise different facets of the same dataset: their approach highlights immune-related female-biased signatures after more extensive covariate adjustment, while ours is more sensitive to modest male-biased and Y-linked signals retained under raw-count modelling. Additionally, our study extends their work by providing a stage-resolved, variance-guided framework that harmonises early and late gestational analyses and incorporates pathway-level and vascular-focused enrichment, offering a broader view of how sex-biased transcriptomic architecture evolves across pregnancy. Analyses of the same dataset can highlight different biological patterns because computational workflows handle covariates, low-abundance genes, and technical replicates differently. These differences reflect the assumptions each framework makes regarding which variation to retain or adjust for, without altering the underlying data. For instance, including birthweight as a covariate removes expression differences linked to fetal growth, whereas excluding it retains those signals. FPKM filtering down-weights low-abundance transcripts, including Y-linked genes, while raw-count models preserve them. Modelling sequencing lanes independently versus collapsing lane-level counts captures different technical variation. When interpreted together, the analyses from our study and that of Olney et al ^26^ provide concordant and complementary views of the same dataset, converging on strong sex-chromosome effects and female-enriched immune signatures, while highlighting different autosomal components of term placental biology under distinct modelling assumptions.

Our study has several limitations that should be acknowledged. First, despite the relatively large sample sizes of both cohorts in placental transcriptomics, bulk RNA-seq averages signals across heterogeneous cell populations and cannot determine whether observed sex-biased differences reflect alterations in cell-type composition, cell-intrinsic transcriptional programs, or both. Second, while we adjusted for key technical and biological covariates, certain factors (e.g., mode of conception, detailed clinical metadata) were either collinear with sequencing batch or too imbalanced to include, potentially introducing residual confounding. Third, the autosome-only analysis at term yielded single DEG under our predefined statistical thresholds, limiting downstream interpretation and restricting pathway analysis to the whole-genome dataset at that developmental stage. Fourth, although preprocessing and normalization pipelines were harmonized across cohorts, the first and term datasets originated from distinct studies, and residual inter-cohort technical variation remains a possibility despite the consistent modelling frameworks employed. Fifth, several maternal covariates known to influence placental transcription (e.g., BMI, age, parity, inflammatory status, medication exposure) were unavailable or incomplete, thereby limiting our ability to adjust for maternal influences on sex-biased expression. Sixth, pathway enrichment analyses rely on annotation databases whose redundancy, granularity, and curation depth may affect the identified pathways. Finally, our analysis is transcriptional and cross-sectional; therefore, we cannot directly infer protein-level activity, cellular localization, or causal mechanisms underlying sex-specific placental differences. Future single-cell and spatial transcriptomic studies will be essential for determining whether the sex-biased signatures we observe arise from shifts in cell-type composition, cell-state differences within shared lineages, or both. By resolving transcription at cellular and spatial resolution, these approaches can reveal sex-specific regulatory programs within trophoblast, endothelial, and immune subpopulations, which are obscured in bulk data ^60,61^.

Despite these constraints, the present study offers several important contributions. Through the analyses of two independent cohorts spanning early and late gestation within fully harmonized preprocessing, modelling, and enrichment frameworks, we provide one of the most comprehensive stage-resolved assessments of placental sexual dimorphism to date, while identifying early placental vascular programming as a key axis of difference. Notably, our targeted vascular analyses uncover that placentas from male fetuses in early gestation exhibit pronounced angiogenesis, matrix remodelling, and hemogenic support, whereas placentas from female fetuses at term display selective engagement of vascular–immune integration and barrier-maintenance pathways. These patterns are not detectable through DEG-level analysis alone. The male-biased placental strategy favoring rapid vascular expansion aligns with established male-biased fetal growth but appears less redundantly organized for immune buffering, limiting its capacity to contain maternal inflammatory signals and maintain effective perfusion and barrier function. In contrast, female-biased transcriptional patterns may represent adaptations supporting fetal homeostasis and could contribute to reduced susceptibility to early-life inflammatory exposures, which have been epidemiologically linked to sex-biased neurodevelopmental outcomes ^47^. Viewed within the developmental origins of health and disease framework, these sex-specific vascular programs may contribute to understanding perinatal outcomes and well-documented sex-biased risks for morbidity, neurodevelopmental issues, and later-life disease, providing an evolutionary perspective linking placental sex with differential vulnerability to inflammation.

In sum, this work demonstrates that sex differences in the human placenta arise from a constellation of transcriptomic processes whose relative contributions reconfigure across gestation, instead of a single static signature. Integrating two developmental windows within a unified analytical framework revealed that sex-biased transcription is structured, multi-layered, and developmentally timed, hallmarks easily overlooked in single-stage analyses. Our findings provide mechanistic insights for sex-specific patterns in fetal growth trajectories, stress adaptation, and pregnancy complications, positioning vascular-immune interfaces as future targets for cell-type-resolved studies to refine models of placental physiology and maternal-fetal health.

## Acknowledgments

We thank Tiago Figueiredo for creating artwork used in the graphical abstract and the Figures 1 (https://tiagofigueiredo.myportfolio.com/work) and Professor Melissa A. Wilson, Senior Investigator at the National Human Genome Research Institute, National Institutes of Health, for kindly providing the term placenta datasets.

## Supplementary Figures and Tables

**Supplementary Table 1:**
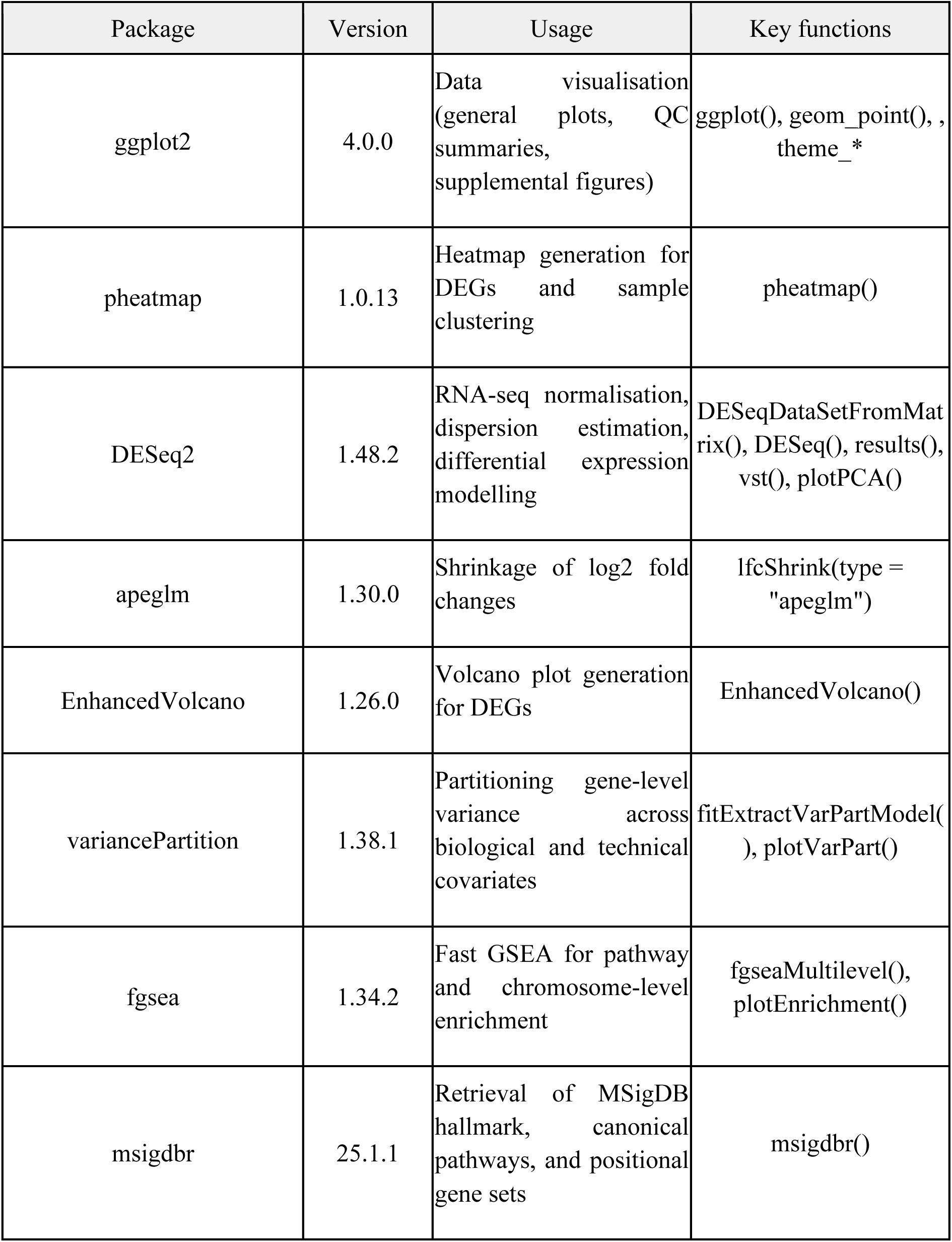
Major Packages used and versions.

### Quality control assessment

**Supplemental Figure 1:**
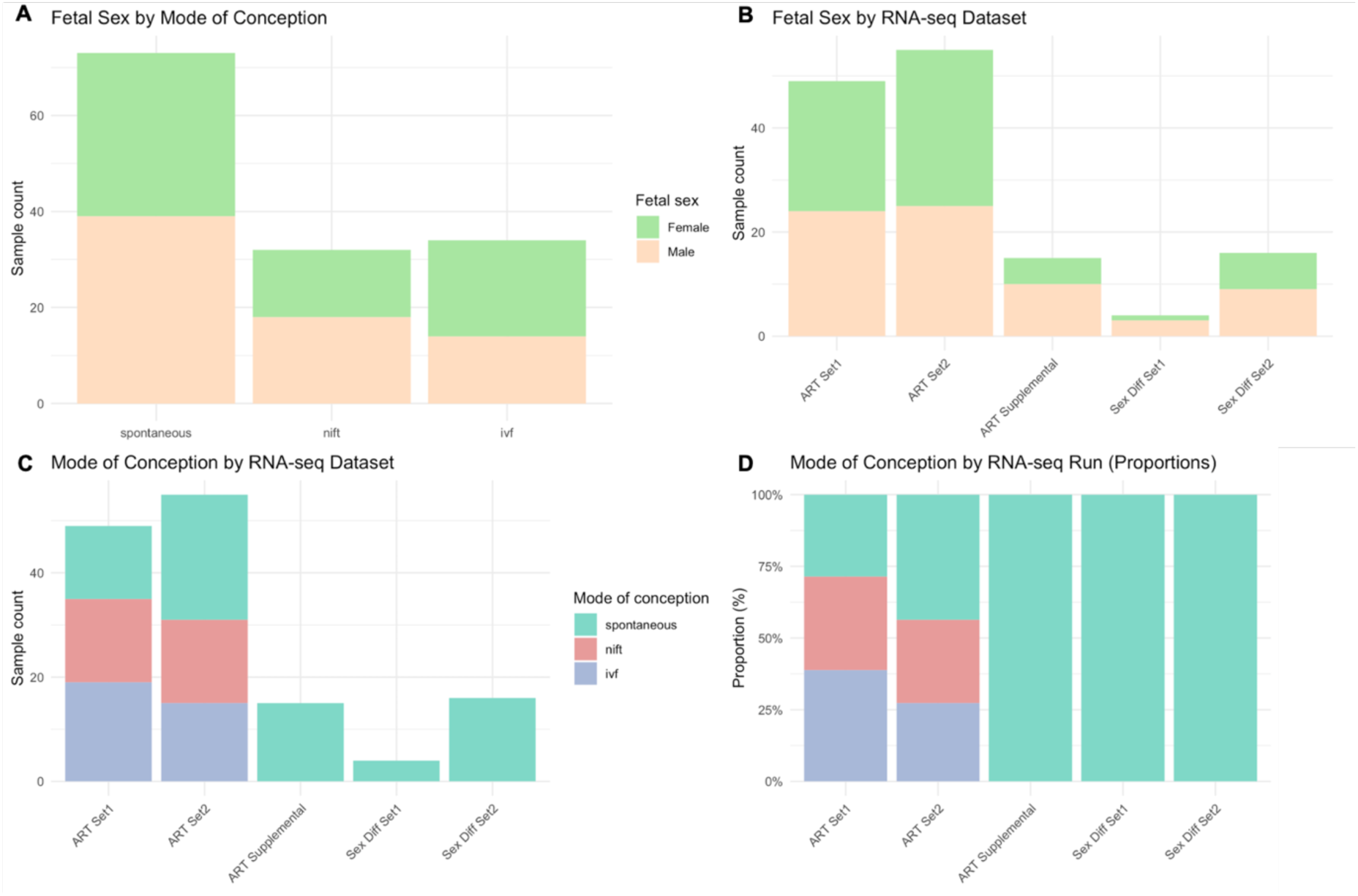
Visualization of sample distributions for the first trimester placenta database. A) Distribution of *fetal sex* by *mode of conception*. B) Distribution of *fetal sex* by *RNA-seq dataset*, C) Distribution of *mode of conception* by *RNA-seq dataset* and D) Proportional distribution of *mode of conception* by *RNA-seq run*.

**Supplemental Figure 2:**
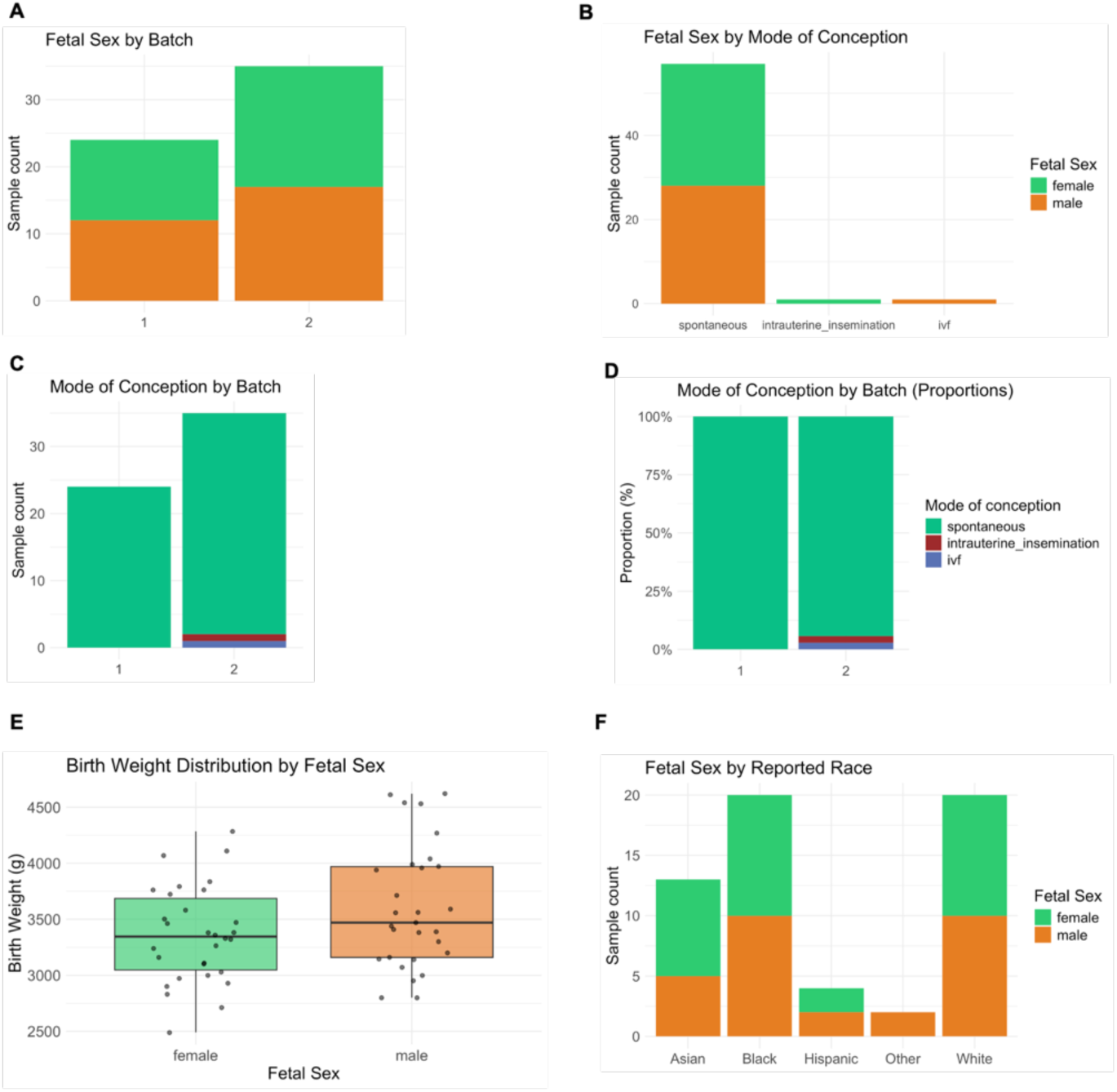
Sample Distribution Visualizations for the Term Placenta Database. A) Distribution of *sex* by *batch*. B) Distribution of *sex by mode of conception*. C) Distribution of *mode of conception* by *batch*. D) Proportional distribution of *mode of conception* by *batch*. E) Boxplot of *birth weight* by sex. F) Distribution of *sex* by *reported race*.

### Variance partition

**Supplemental Figure 3:**
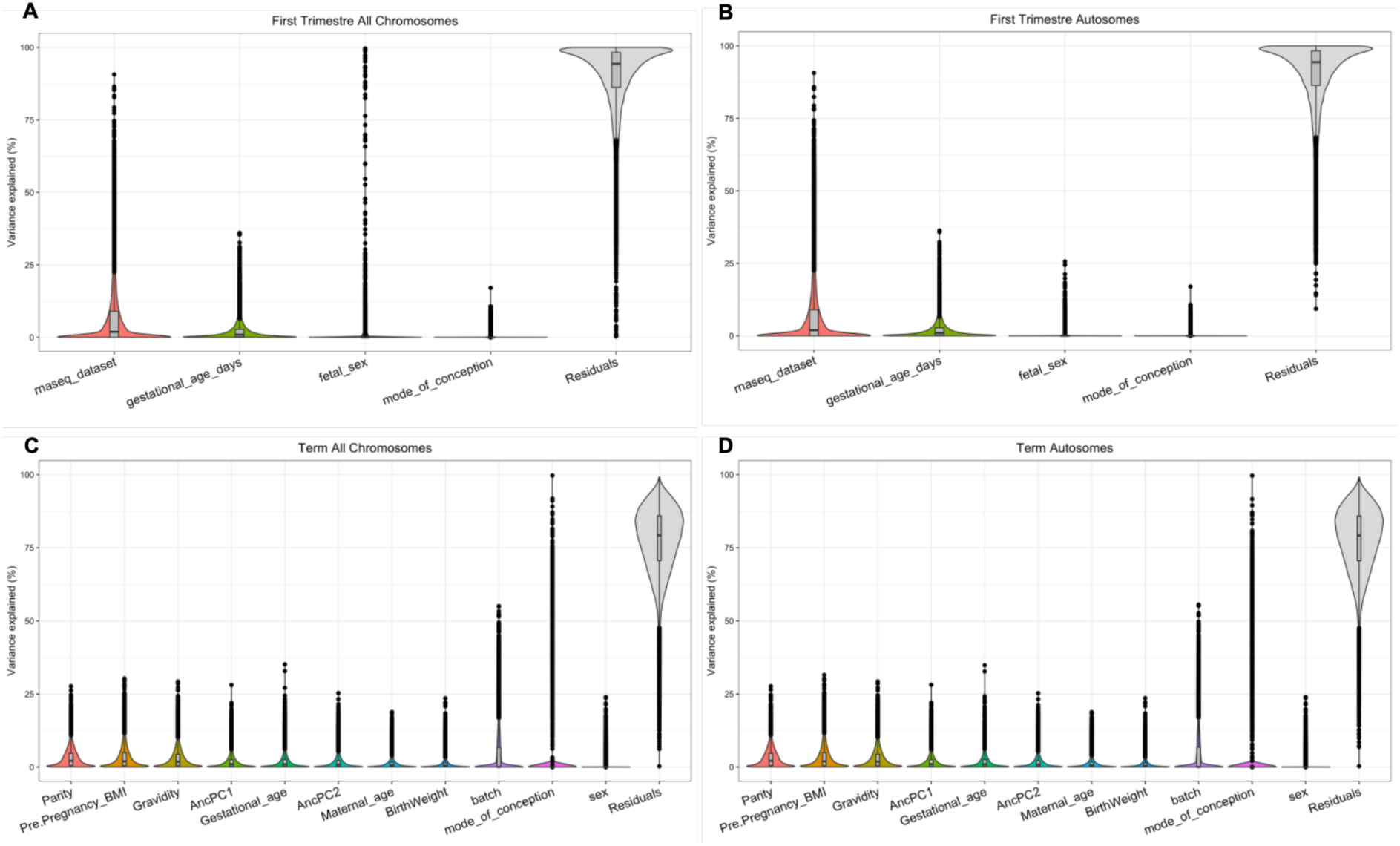
Variance Partition Analysis of First Trimester and Term Placental Gene Expression. Violin plots illustrate the distribution of variance explained by each study variable across all genes, with median and quartile values displayed as overlaid box plots. Variance partition analysis was performed to identify variables contributing meaningfully to gene expression variation and to inform DESeq2 model specification. A) First Trimester, All Chromosomes; B) First Trimester, Autosomes Only; C) Term Placenta, All Chromosomes; D) Term Placenta, Autosomes Only.

**Supplemental Figure 4:**
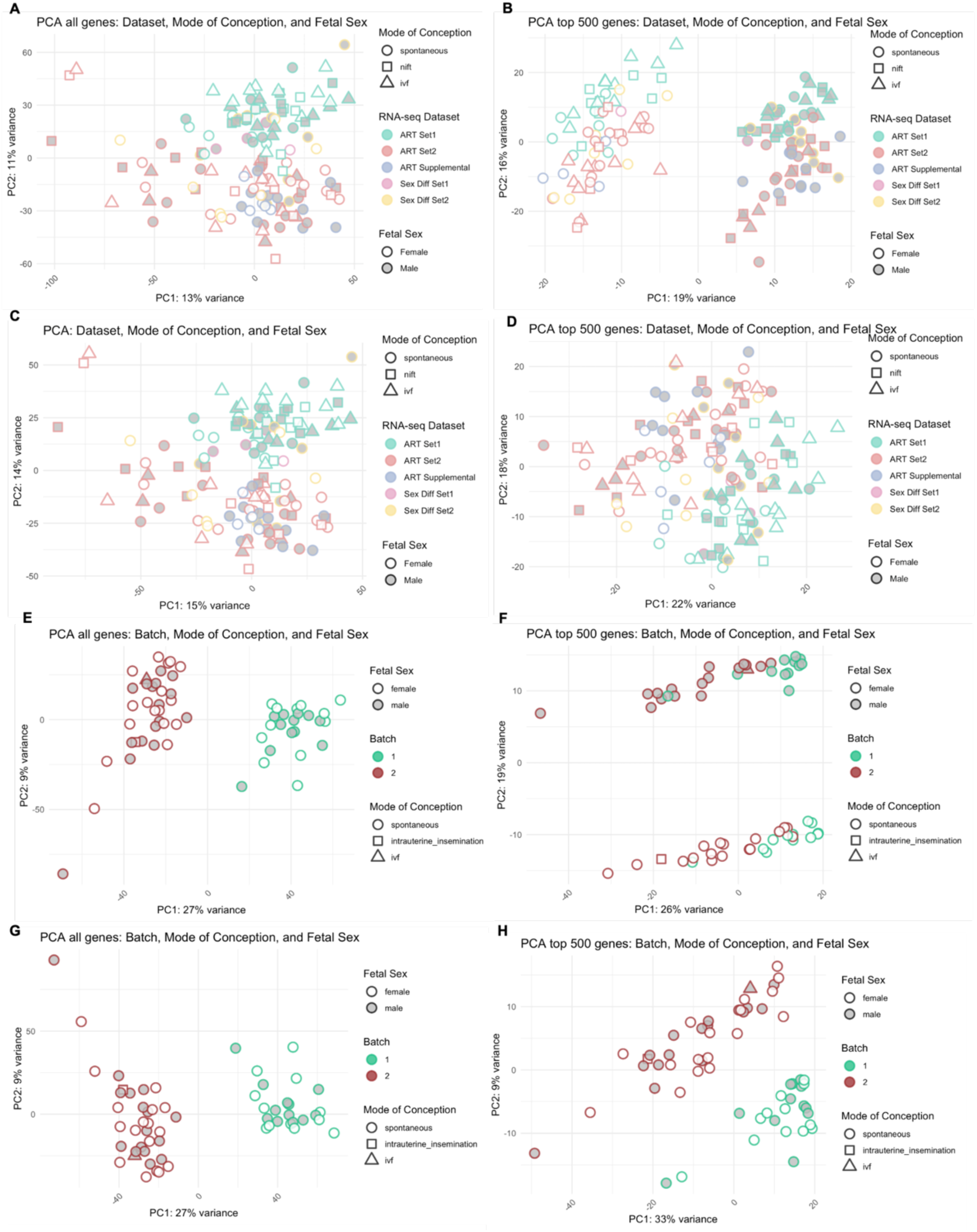
PCA plots illustrating gene expression data from first trimester and term placenta databases. Plots were generated using either the entire gene set or the top 500 genes, for both the whole genome and autosomal subsets. A-D) First trimester placenta database, clustered according to *dataset*, *mode of conceptio*n and *fetal sex*: A) Whole genome (all genes); B) Whole genome (top 500 genes); C) Autosomes (all genes); D) Autosomes (top 500 genes). E-H) Term placenta dataset, clustered according to *dataset*, *mode of conception* and *fetal sex*: E) Whole genome (all genes); F) Whole genome (top 500 genes); G) Autosomes (all genes); H) Autosomes (top 500 genes).

### Rank-Based GSEA (FGSEA)

**Supplemental Figure 5:**
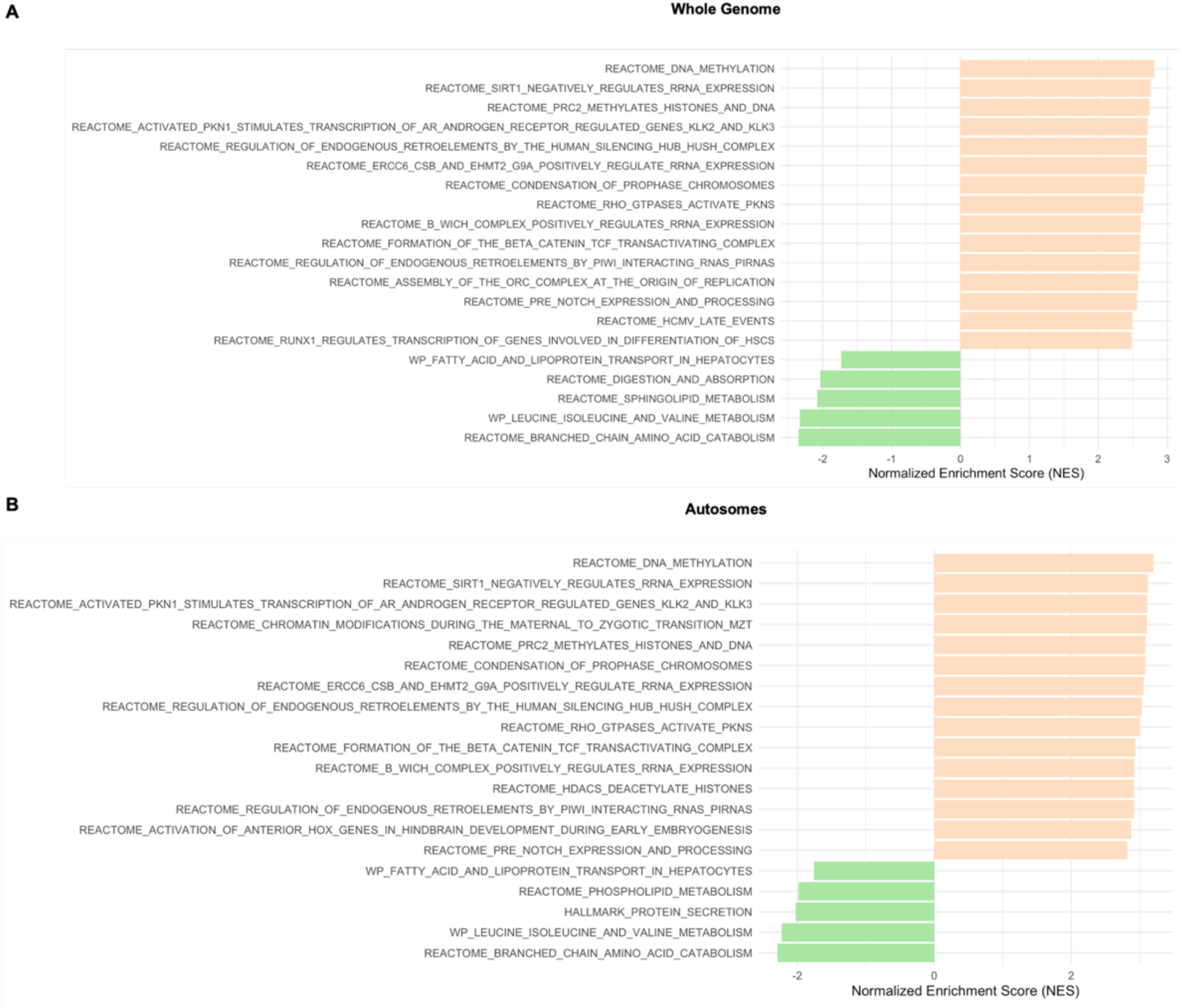
Rank-based gene set enrichment analysis (fgsea) of fetal sex-biased gene expression in first trimester placenta. A) Whole genome and B) Autosomal genes only. Bars represent the 20 most significantly enriched pathways (FDR < 0.05), ranked by normalized enrichment score (NES). Positive NES values (orange) indicate enrichment in male placentas; negative NES values (green) indicate enrichment in female placentas. Pathway databases queried: Hallmark (H), KEGG (C2:CP:KEGG_MEDICUS), Reactome (C2:CP:REACTOME), WikiPathways (C2:CP:WIKIPATHWAYS), BioCarta (C2:CP:BIOCARTA), and Pathway Interaction Database (C2:CP:PID) from the Molecular Signatures Database (MSigDB v25.1.1).

**Supplemental Figure 6:**
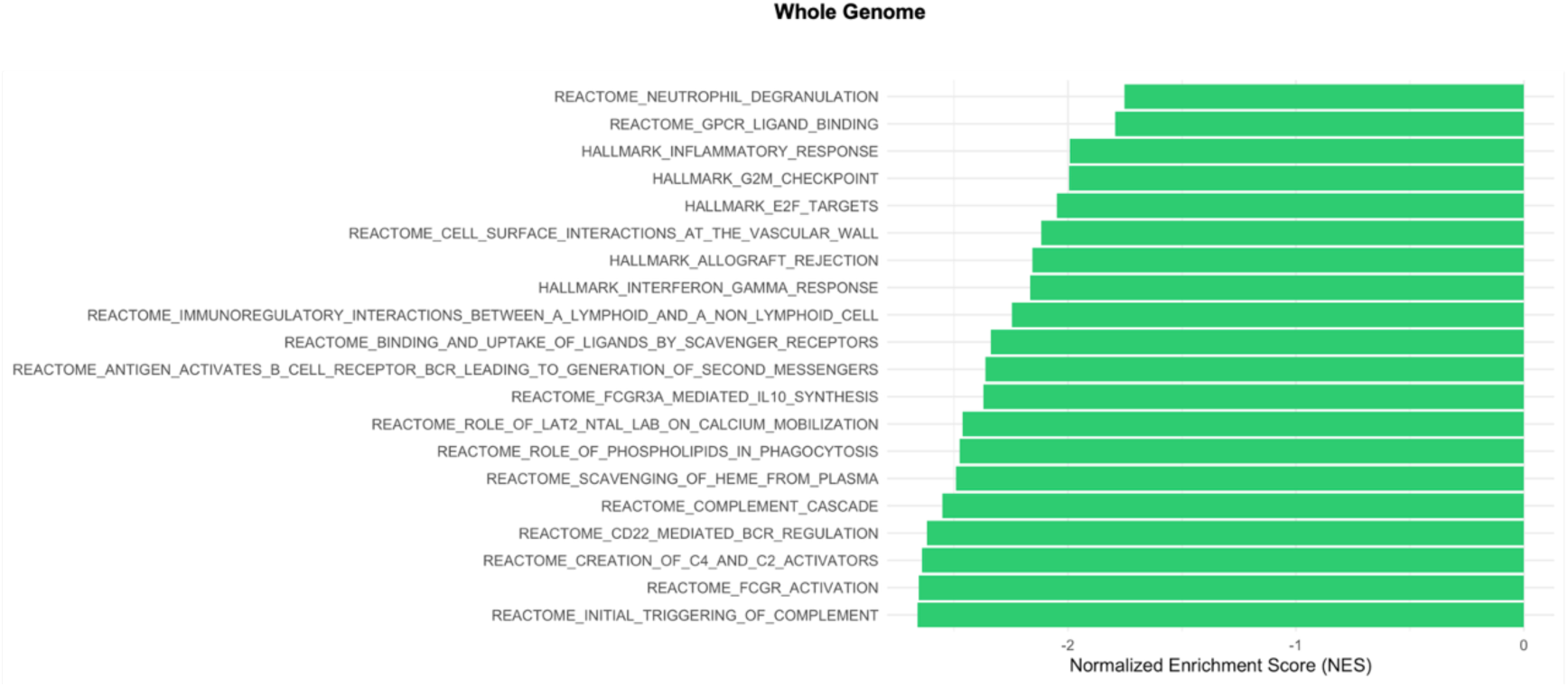
Rank-based gene set enrichment analysis (GSEA) of fetal sex-biased gene expression in term placenta (whole genome). Bars represent the 20 most significantly enriched pathways (FDR < 0.05) ranked by normalized enrichment score (NES). Negative NES values (green) indicate enrichment in female placentas. Pathway databases queried: Hallmark (H), KEGG (C2:CP:KEGG_MEDICUS), Reactome (C2:CP:REACTOME), WikiPathways (C2:CP:WIKIPATHWAYS), BioCarta (C2:CP:BIOCARTA), and Pathway Interaction Database (C2:CP:PID) from the Molecular Signatures Database (MSigDB v25.1.1).

### FGSEA chromosome analysis

**Supplemental Figure 7:**
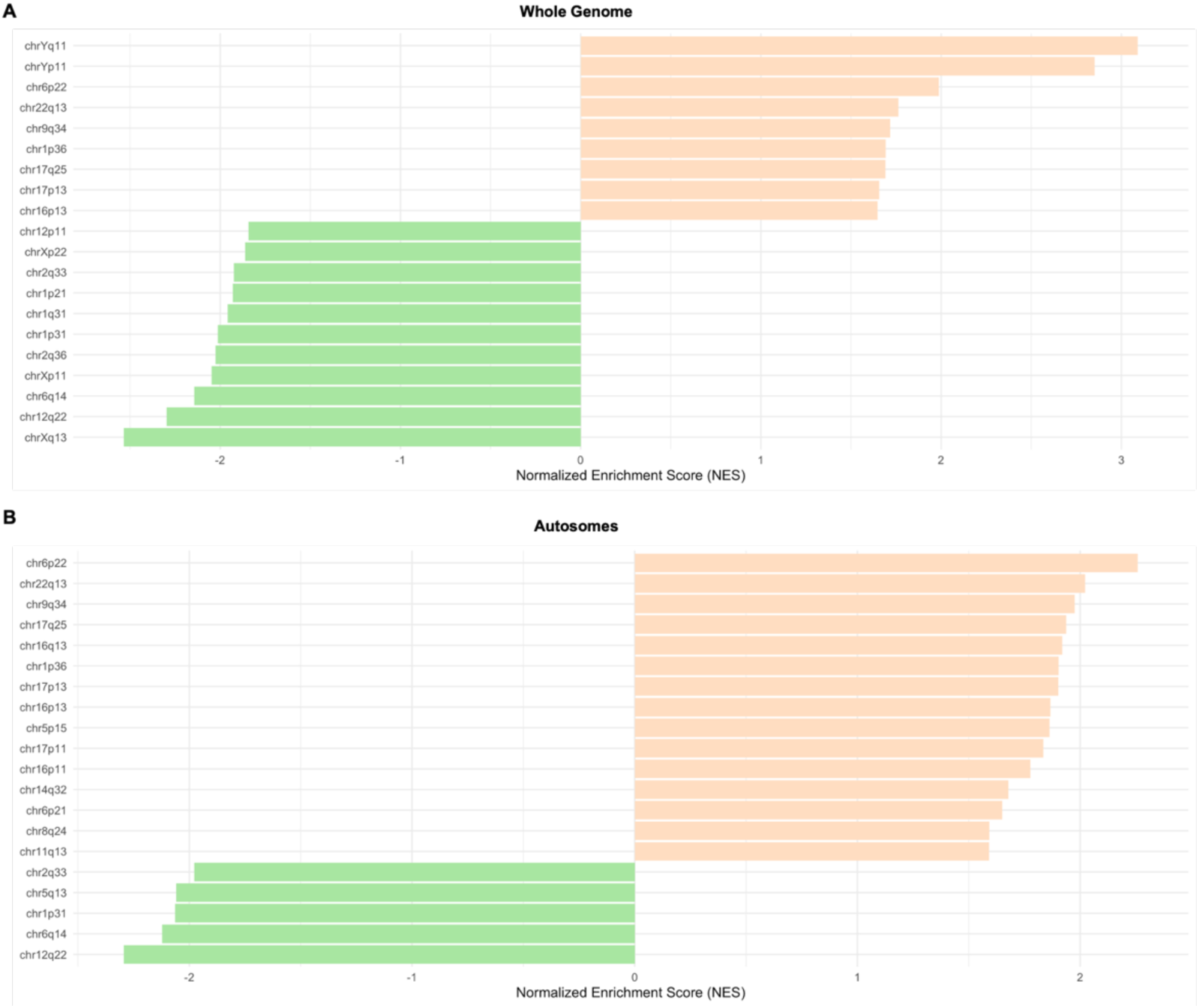
Chromosome-based gene set enrichment analysis (GSEA) in first trimester placenta. A) Whole genome and B) Autosomal genes only. Bars represent the 20 most significantly enriched chromosome position gene sets (FDR < 0.05) ranked by normalized enrichment score (NES). Positive NES values (orange) indicate enrichment in male placentas; negative NES values (green) indicate enrichment in female placentas. Chromosome gene sets are from the C1 collection of the Molecular Signatures Database (MSigDB v25.1.1).

**Supplemental Figure 8:**
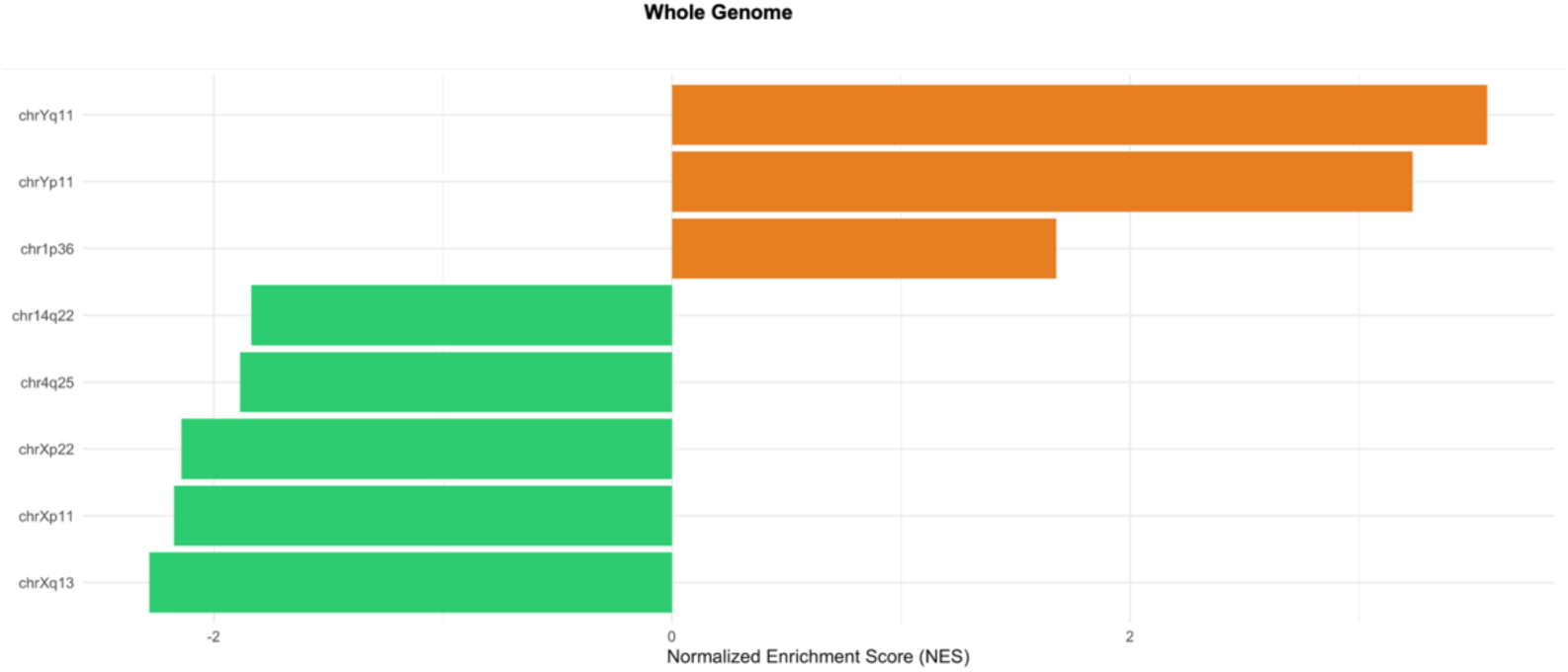
Chromosome-based gene set enrichment analysis (GSEA) was performed on term placentas (whole genome). Bars represent the 20 most significantly enriched chromosome position gene sets (FDR < 0.05), ranked by normalized enrichment score (NES). Positive NES values (orange) indicate enrichment in male placentas; negative NES values (green) indicate enrichment in female placentas. Chromosome gene sets are from the C1 collection of the Molecular Signatures Database (MSigDB v25.1.1).

